# Parasite manipulation of the host cell cycle as a means to block inflammatory signaling and promote intracellular replication

**DOI:** 10.1101/625194

**Authors:** Zhee Sheen Wong, Sarah L. Sokol, J. P. Dubey, Jon P. Boyle

**Affiliations:** Department of Biological Sciences, Dietrich School of Arts and Sciences, University of Pittsburgh, Pittsburgh, Pennsylvania, United States of America; Animal Parasitic Diseases Laboratory, Beltsville Agricultural Research Center, Agricultural Research Service, U.S. Department of Agriculture, Beltsville, Maryland, United States of America

## Abstract

*Toxoplasma gondii* and *Hammondia hammondi* are closely-related coccidian intracellular parasites that differ in their ability to cause disease in animal and (likely) humans. The role of the host response in these phenotypic differences is not known and to address this we performed a transcriptomic analysis of a monocyte cell line (THP-1) infected with these two parasite species. The pathways altered by infection were shared between species ~95% the time, but the magnitude of the host response to *H. hammondi* was significantly higher compared to *T. gondii*. Accompanying this divergent host response was an equally divergent impact on the cell cycle of the host cell. In contrast to *T. gondii*, *H. hammondi* infection induces cell cycle arrest via pathways linked to DNA-damage responses and cellular senescence and robust secretion of multiple chemokines that are known to be a part of the senescence associated secretory phenotype (SASP). Remarkably *T. gondii*-conditioned media can suppress the SASP response during *H. hammondi* infection, and this suppression is accompanied by a corresponding increase in the replication rate of *H. hammondi* in recipient cells. Taken together our data suggest that *T. gondii* manipulation of the host cell cycle provides a novel mechanism to avoid stress and/or DNA-damage induced responses by the host cell, and that this ability has a direct impact on parasite replication rate both within the host cell as well as in bystander cells.

## Introduction

The complex interaction between the intracellular pathogen *Toxoplasma gondii* and its host cell can lead to extensive changes in the host transcriptome [1–3]. These changes can have dramatic effects on host phenotypes including those responsible for the regulation of metabolism, apoptosis and innate immune signaling [4–6]. A successful infection in *T. gondii* results in rapid replication of infecting parasites as tachyzoites, dissemination to a variety of tissues, followed by the clearance of actively replicating stages and encystment of bradyzoite stages in muscle and neuronal tissues. Therefore a balance exists for *T. gondii* to expand its numbers during the acute phase such that a sufficient number of parasites can make it to the chronic, cyst stage and therefore be transmitted to the next host.

It has long been known that *T. gondii* infection dramatically alters the transcriptional landscape in the host cell [4], and many of these changes can be directly linked to the secretion of specific effectors directly into the host cell [7,8], and *T. gondii* is likely to harbor hundreds of such effectors [9–12]. Rhoptry-derived proteins such as ROP16 regulate important transcription factors such as STAT3 [13] and STAT6 [14], while dense granule proteins such as GRA15 [15], GRA16 [16] and GRA24 [17] modulate host NF-κB, p53 and p38 MAP kinase host pathways, respectively. GRA18 has recently been shown to modulate the Wnt signaling pathway in mouse cells, owing to its ability to stabilize the transcription factor β-catenin [18], and GRA25 induces CCL2 in human foreskin fibroblasts [19]. In addition, parasite effector TgIST represses the interferon (IFN)-γ response by recruiting the Mi-2/NuRD repressor complex and subsequently blocking STAT1-related IFNγ-stimulated transcription [20,21]. Many dense granule effectors including GRA16, GRA18 and GRA24 require the presence of a protein complex on the parasitophorous vacuole membrane (PVM) consisting of at least three proteins named MYR1, 2 and 3 [22,23]. This complex has also been shown recently to be critical for the induction of the chemokine CCL22 in human placental cells [24]. Importantly *T. gondii* has no impact on CCL22 production by human foreskin fibroblasts (HFFs) [24], further demonstrating that responses to *T. gondii* infection can vary dramatically across cell types. Taken together it is clear that multiple means of modulating the host cell environment have evolved in *T. gondii* and the sum total of these effects is an essential component of its compatibility with its many hosts. *Hammondia hammondi* is the closest extant relative of *T. gondii*, and was first discovered in the feces of a cat in Iowa, USA in 1975 [25]. Unlike *T. gondii, H. hammondi* has a restricted natural intermediate host range, having been known to infect rats, mice, goat and roe deer in the wild [26–28], and in the laboratory non-human primates can be infected but not birds [26–30]. *H. hammondi* shares >99% of its ~8000 genes in near perfect synteny with *T. gondii* and expresses functional orthologs of key *T. gondii* effectors such as ROP18 and ROP5 [31–33]. This is important from an epidemiological and evolutionary perspective, as *H. hammondi* is not known to cause clinical disease in any naturally infected intermediate (human and other animals) or definitive host [33]. In addition, in tissue culture *H. hammondi* replicates for a short period of time and then enters into a unique terminally differentiated bradyzoite state that is a) unable to be subcultured *in vitro*, b) incapable of infecting an intermediate mouse host and c) only capable of infecting a feline host [29]. Paradoxically, despite this natural (and possible pre-determined) bradyzoite developmental program and expression of canonical bradyzoite genes during tachyzoite-like replication, *H. hammondi* cannot be induced to form bradyzoites using stresses like high pH medium that robustly induce cyst formation in *T. gondii* [34]. While it is not yet known if *H. hammondi* is capable of infecting humans (and there are no tests as yet capable of distinguishing these closely-related species serologically), *H. hammondi* and *T. gondii* oocysts are known to co-circulate, suggesting that humans may encounter this parasite [34,35].

Since multiple *T. gondii* effectors modulate host transcriptional regulation, and to date many of these have been found to be functionally conserved between *T. gondii* and *H. hammondi* [31,36], we sought to test the hypothesis that differences in the response of the host cell to *T. gondii* and *H. hammondi* could be linked to phenotypic differences between these species, including replication rate and pathogenesis. To do this we performed the first thorough analysis of the cell-autonomous host response to *H. hammondi* and compared it to *T. gondii*. We found that the majority of infection-induced changes in the host cell were conserved between these two species, but also that the magnitude of these changes in *H. hammondi*-infected cells was much larger (often by many-fold) than in *T. gondii*-infected cells. We confirmed that this effect is manifested at the protein level for a subset of chemokines, and that a similar effect could be observed *in vivo* during mouse infections. In addition, we also identified a small, but important, subset of host cell pathways that were altered during *T. gondii* infection but were either unaltered or even inverted in *H. hammondi*-infected cells. The central theme in these differentially regulated pathways was a role in the host cell cycle, which *T. gondii* is well known to manipulate [37,38]. These data and follow up experiments described here suggest that a) *T. gondii* and *H. hammondi* have distinct effects on the host cell cycle, b) *T. gondii* infected cells produce significantly less chemokine in response to infection compared to *H. hammondi* infected cells, c) *H. hammondi*-infected cells closely resemble cells undergoing senescence and display something akin to a senescence-associated secretory phenotype (SASP), and d) all of these effects serve to increase the replication rate of *T. gondii*. Overall this work has linked cell cycle manipulation by *T. gondii* as a global mechanism of transcriptional suppression, and determined that this is yet another unique adaptation of *T. gondii* compared to its near relatives. Remarkably, *T. gondii* infection leads to paracrine signals that markedly increase the sensitivity of neighboring cells to parasite infection.

## Results

### *H. hammondi* sporozoites induce a more potent differential gene expression in THP-1 cells as compared to *T. gondii* during acute infection

To compare the impact of *T. gondii* and *H. hammondi* infection on the transcriptional host response during infection, we performed RNA-sequencing (seq) on THP-1 cells (a monocyte cell line) infected with representatives of three major lineages of *T. gondii* and two *H. hammondi* isolates. In all cases (unless specified), we used parasites excysted from oocysts that were grown for 24 hours on HFFs.

We first used principal component analysis (PCA) to look broadly at differences and similarities in the host transcriptome across strains and species [39]. According to this analysis the first PC (PC1, 68% variance) indicated a clear separation between parasite species, while the second (PC2, 20% variance) encompassed differences between strains of each species. These data indicate that THP-1 cells infected with all three *T. gondii* types (TgGT1, TgME49 and TgVEG) are more similar to each other than to THP-1 cells infected with either strain of *H. hammondi* (HhEth1 and HhAmer; Fig. 1A). Infection status (Mock vs. Infected) also was distributed across PC1, in that mock-infected cells clustered together within PC1 regardless of strain or species (Fig. 1A). Importantly the two *H. hammondi* strains were much further separated from all mock-infected samples along this PC, suggesting 1) a distinct response by the host cells to this species and/or 2) more potent induction of transcriptional changes by the parasite. PC2 showed a clear separation between TgGT1-infected cells and all other samples, including *T. gondii* and *H. hammondi*. While it was clear from PCA and other analyses (below) that TgGT1 infection induced extensive changes in transcript abundance compared to mock-infection (Figs. 1A, C and H), the TgGT1-infected cells were clearly the most distinct *T. gondii* strain along PC2. Therefore we have excluded TgGT1 infection in some of the downstream analysis. Hierarchical clustering of the distances between samples also further confirmed the differential expression profiles seen in THP-1 cells infected with *H. hammondi* as compared to *T. gondii* (Fig. 1B).

**Fig. 1.**
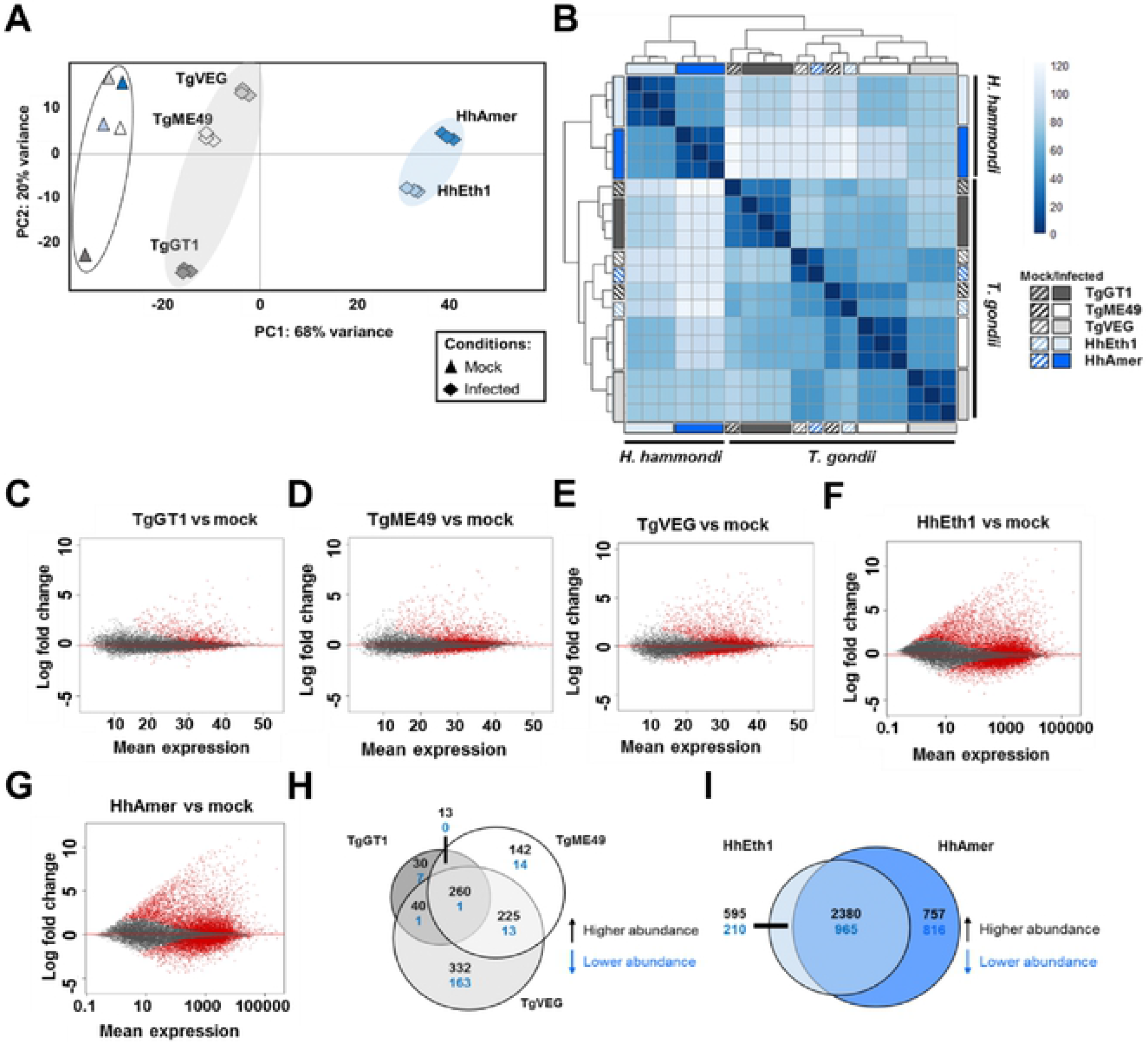
Unique transcriptome profiles of THP-1 cells infected with parasites where *H. hammondi* induced more dramatic numbers of differentially expressed genes in THP-1 cells than *T. gondii.* (A) Principal components(PC) 1 and 2 of THP-1 cells infected(♦) or mock infected(▴) with *T. gondii*(TgGT1 (Dark grey), TgME49(white) or TgVEG(light grey)) or *H. hammondi*(HhEth1 (light blue) or HhAmer(dark blue)). THP-1 cells infected with *T. gondii* clustered together(light grey shading) whilst THP-1 cells infected with *H. hammondi* clustered together along PC1 (light blue shading). *T. gondii* and H. hammondi mock-infected cells(white shading) also separated from the parasite­ infected cells.(B) Heatmap showing Euclidean distances clustering between mock (striped boxes) and infected(solid filled boxes) THP-1 cells samples(log_2_-transformed data).(C-G) MA-plots of genes expression in THP-1 cells infected with T gondii(Tg) or *H. hammondi*(Hh). Red dots 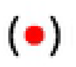 represent genes of different abundance in infected THP-1 cells as compared to mock-infected THP-1 cells(alpha= 0.01).(H-1) Venn diagrams of genes in THP-1 cells infected with *T. gondii*(H) or *H. hammondi*(I; *P*_*adj*_ < 0.01, log_2_ fold-change ≥ 1 or ≤−1).

We compared host gene expression of parasite-infected cells to mock-infected cells for all types/strains using *DESeq2* [40] with the thresholds of log_2_ fold-change ≥ 1 or ≤ −1 with *p*_*adjuster (adj)*_ < 0.01 and identified host transcripts that were of higher or lower abundance in response to infection (Table S1). The most dramatic outcome of this experiment was the striking difference in the host transcriptional response to *H. hammondi* compared to *T. gondii*, regardless of strain. Specifically, HhEth1 and HhAmer-infected THP-1 cells had significant changes in ~27% and ~32% of the 15452 queried transcripts, respectively (Fig. 1I and Table S1), while in *T. gondii* this ranged from ~4% for TgGT1 to ~7% for TgVEG (Fig. 1H and Table S1). MA-plots showing average expression vs. log fold-change of each gene, further illustrate the stark contrast in the extent of changes in transcript abundance that are induced by *T. gondii* and *H. hammondi* (Figs. 1C-E vs. F-G). When we compared sets of differentially expressed genes across strains of each species, we identified type and/or strain-specific gene expression profiles (Figs. 1H and I). It is also interesting to note that more strain type-specific changes in transcript abundance were observed among the *T. gondii* strains compared to HhEth1 and HhAmer infections (Figs. 1H vs. I).

### Hyperinduction of chemokines by *H. hammondi* compared to *T. gondii* is recapitulated *in vivo*

While differences in host responses by *T. gondii* and *H. hammondi* were readily apparent *in vitro*, little is known about differences in the host response to these parasites *in vivo*. Therefore we infected mice intraperitoneally with parasites of each species and collected peritoneal lavage supernatants and peritoneal cells. Some of the highly differentially expressed genes by *T. gondii* and *H. hammondi in vitro* infection were the host cytokine genes (Table S1). Therefore we analyzed cytokine transcripts by reverse transcriptase (RT)-qPCR and measured cytokine levels in the supernatants by ELISA (Fig. 2). Since Il12 and Ifnγ are cytokines that were induced in response to *T. gondii* infection and Il12 plays an important role in activating Ifnγ during *T. gondii* infection [41–43], we quantified mouse Il12p40 and Ifnγ in peritoneal cell supernatants taken from mice infected with TgVEG or HhAmer at various time points to ensure that mice were infected (Figs. 2SF and G). Since *H. hammondi* replicates ~4X slower than *T. gondii in vitro* [34], we also monitored relative replication rates and parasite loads using species-specific RT-qPCR for the parasite *GRA1* gene. Our data from the RT-qPCR showed that at any given time point, *H. hammondi* parasite burden was significantly lower than *T. gondii* (Sidak’s multiple comparisons test, \*\**p*<0.01, \*\*\**p*<0.001 and \*\*\*\**p*<0.0001 as compared to the respective time points), indicating that *H. hammondi* also replicates more slowly than *T. gondii in vivo* (Fig. 2SE). We also observed that *H. hammondi* sporozoites have a lower viability post-excystation as evidence by *GRA1* levels at 30 min post-infection (Fig. 2SE; Sidak’s multiple comparisons test \*\**p*<0.01, \*\*\**p*<0.001 and \*\*\*\**p*<0.0001).

**Fig. 2.**
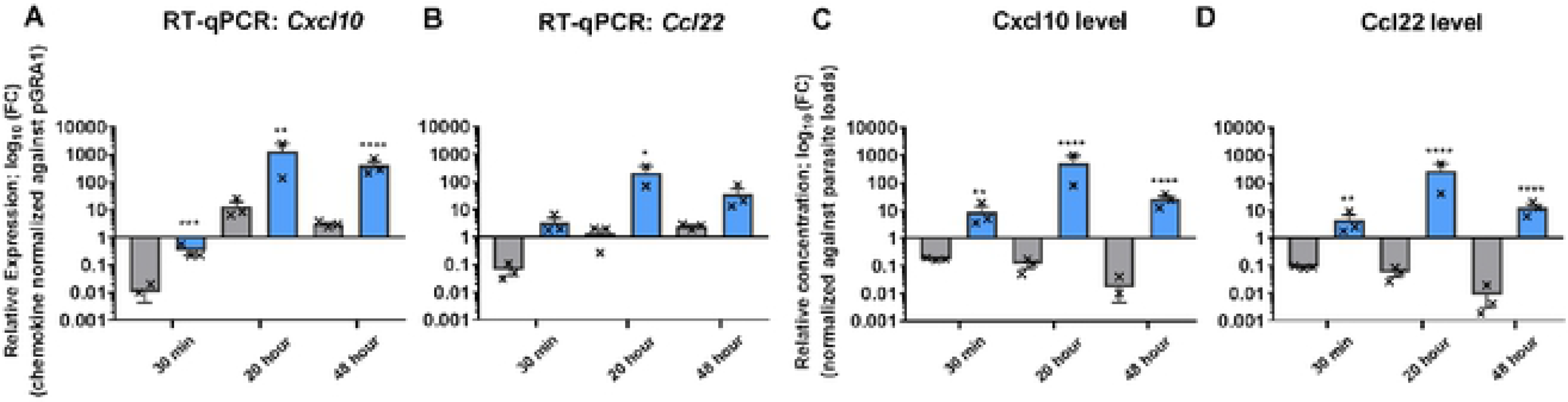
*In vivo H. hammondi* infection leads to higher levels of chemokine transcripts and protein compared to T. *gondii* when parasite burden is taken into account. Mice were infected with *T. gondii* (TgVEG; grey) or *H. hammondi* (HhAmer; blue). Mice were also mock-infected with parasite free extracts. Mouse peritoneal cell RNA and supernatants were collected at 30 min, 20 and 48 h post sporozoite infections. mRNA levels of chemokine genes were quantified using RT-qPCR and Gapdh was used as the reference gene. Parasite GRA1 was used as the reference when parasite burden was taken into account. Protein levels were quantified using ELISA. (A-B) Bar graphs show relative expression (RE; log_10_) normalized against parasite loads (2^·M^CT calculated from LlCr chemokine genes - ΔC_T_ *GRA* 1). RE of *Cxcl10* was significantly higher in HhAmer-infected mice as compared to TgVEG infection at any time points (Sidak's multiple comparisons test; **p<0.01, ***p<0.001 and ****p<0.0001). RE of *Ccl22* were significantly different in relative to parasite loads at 20 h post-infection (Sidak's multiple comparisons test; *p<0.05). (C-D) Relative concentration of cytokines normalized against parasite loads (2^−ΔΔT^ calculated from log_2_-transformed chemokine protein levels - LlCr *GRA* 1). Cxcl10 and Ccl22 levels were significantly different in response to TgVEG or HhAmer infections at any time point (Tukey's multiple comparisons test, **p<0.01, ****p<0.0001). Error bars represent SEM.

To compensate for this growth difference (which is an intrinsic property of these parasites species), we set up an independent experiment where we normalized cytokine transcript and protein level to *GRA1* transcript level, a reasonable proxy for parasite burden [34]. After infecting mice with 20,000 freshly excysted TgVEG or HhAmer sporozoites, we found that, as expected, *H. hammondi* showed a significantly lower parasite burden over the course of the experiment compared to *T. gondii* (Fig. 2SH, Sidak’s multiple comparisons test *\*\*p*<0.01 and \*\*\*\**p*<0.0001). However, when we normalized cytokine transcript levels (specifically *Cxcl10* and *Ccl22*) in peritoneal cells to parasite numbers we found that *H. hammondi* infection resulted in significantly higher levels of parasite-normalized transcript at 30 min, 20 and 48 h post-infection for *Cxcl10* and 20 h post-infection for *Ccl22* (Figs. 2A and B; Sidak’s multiple comparisons test \**p*<0.05, \*\**p*<0.01, *** *p*<0.001 and **** *p*<0.0001). When we performed the same analysis on parasite burden-normalized Cxcl10 and Ccl22 protein levels in the peritoneal lavage fluid, we also found that *H. hammondi*-infected mice had significantly higher parasite-normalized chemokine levels compared to *T. gondii* (Figs. 2C and D; Tukey’s multiple comparisons test; \*\**p*<0.01and **** *p*<0.0001). These data indicate that the host response to *H. hammondi* is much more robust *in vivo* compared to that in response to *T. gondii* when parasite burden is taken into account.

### *H. hammondi*-infected cells had more robust changes in transcription compared to *T. gondii*, despite extensive overlap in the altered pathways

From PCA and *DESeq2* analysis of THP-1 cells infected with parasites, it is clear that *H. hammondi*-infected cells display a more robust transcriptional change than *T. gondii*-infected cells (Fig. 1). However this could occur by differential induction of distinct sets of genes, and/or via differences in the overall magnitude of the induction of the same gene sets. To assess this we used Pre-ranked Gene Set Enrichment Analysis (GSEA; [44,45]) on each transcriptional profile using the curated “Hallmark” gene sets database. Overall we identified 47 gene sets that were significantly enriched in cells infected with *T. gondii* and/or *H. hammondi* (FDR-q value<0.05), and 41 of these were shared gene sets enriched (positively or negatively) in infected cells compared to mock-treated cells (Figs. 3A and B, Table S2). Interestingly, *H. hammondi* infections induced quantitatively higher enrichment of these shared transcriptional changes. For example, for *IFNγ response* all *T. gondii* and *H. hammondi* strains significantly altered this gene set during infection, but *H. hammondi* Eth1 and Amer had normalized enrichment scores (NES) of 11.1883 and 11.3855 respectively while TgGT1, TgME49 and TgVEG-infected cells had and NES of 8.0227, 8.5538 and 10.0764 respectively (Fig. 3A, Table S2). This is further illustrated by cluster analysis of a subset of the *IFNγ response* gene set which includes multiple *IRF* and chemokine genes (Fig. 3C, Table S3 and Fig. 3Sc (top panels)). Using Ingenuity Pathway Core Analysis (IPA), we also identified 280 canonical pathways significantly enriched in *T. gondii* and/or *H. hammondi* infections (Benjamin-Hochberg adjusted *p* <0.01, Table S4). These pathways include *dendritic cell maturation, IL-1 signaling, IL-6 signaling* and *interferon signaling* (Fig. 3SaA and B). Our IPA analysis reaffirm our GSEA results that while *H. hammondi* infection alters many of the same canonical pathways and gene sets in THP-1 cells as does infection with *T. gondii*, the magnitude of the effect is greater (e.g., *dendritic cell maturation* had *z*-scores of 6.38 and 6.20 for HhAmer and HhEth1 respectively; 5.57, 4.69 and 4.09 for TgGT1, TgME49 and TgVEG respectively; Fig 3SaA, Table S4). This could be due to 1) *T. gondii* suppression of these host responses and/or 2) direct induction of a more dramatic host response during infection by *H. hammondi*. Since IFNγ-signaling is critical for controlling *T. gondii* infection [41–43] we examined all members of the *interferon signaling pathway* and identified a subset of them (*BAK1, 1FITM1, 1FITM2, JAK1, JAK2* and *PSMB8*) that were only differentially expressed in THP-1 cells infected with *H. hammondi* (Fig. 3SbA-C).

**Fig. 3.**
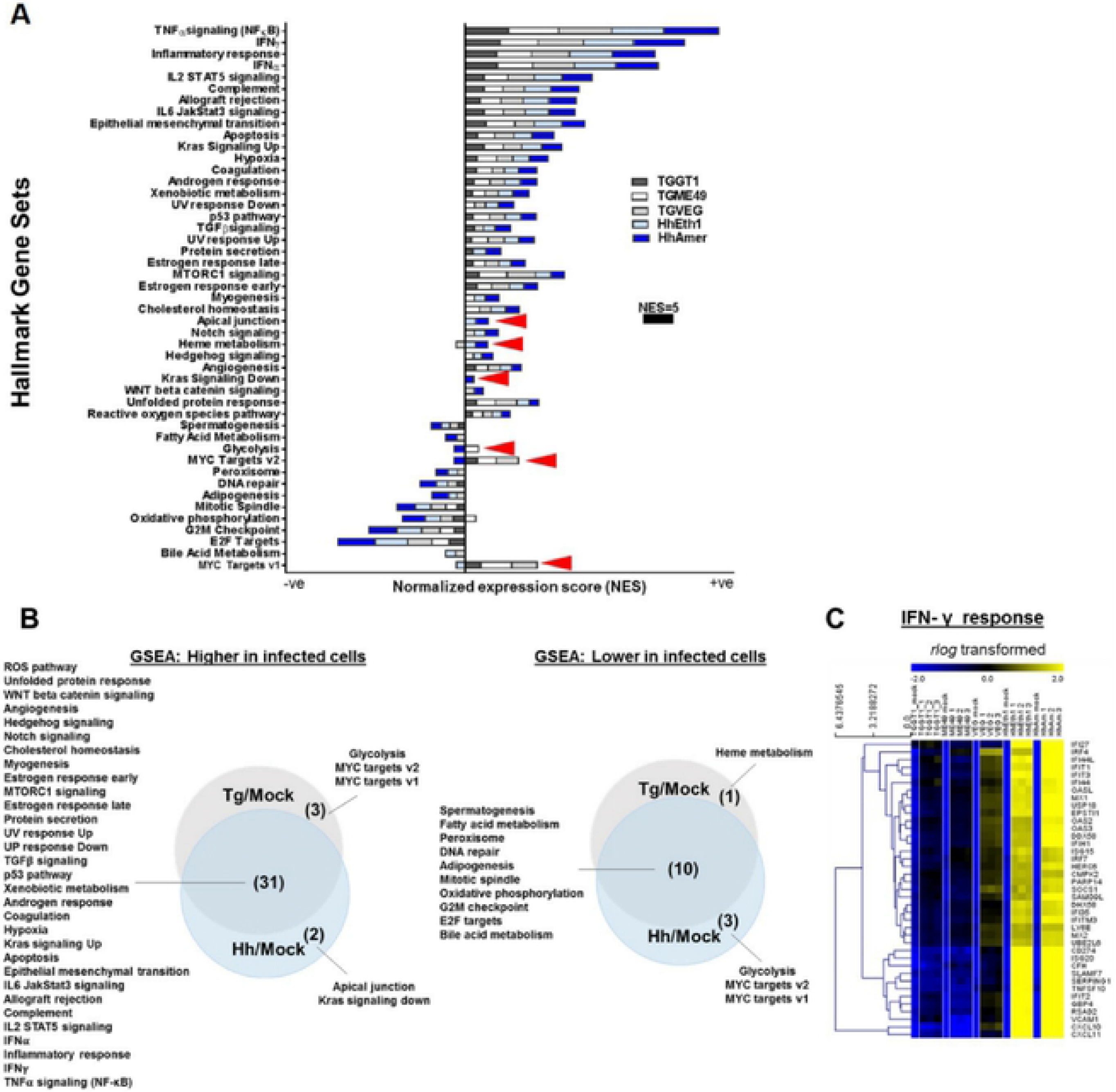
The majority of host pathways altered in *T. gondii* and *H. hammondi-* infected cells are shared between species. *rlog*-normalized data was used to perform Pre-ranked Gene Set Enrichment Analysis (GSEA) to understand biological relevance of the data set. (A) Bar graph shows normalized enrichment scores (NES) of positively (+ve) and negatively (-ve) enriched curated "Hallmark" gene sets in THP-1 cells in response to *T. gondii* (Tg; GT1, ME49 and VEG) and *H. hammondi* (Hh; Eth1 and Amer) infections as compared to mock-infected THP-1 cells. Shown are gene sets with a false discovery rate (FDR-q value) < 0.05 (computed with 1000 Monte-Carlo simulations). The NES represents the degree to which a gene set is overrepresented at the top or bottom of the ranked list. Bars show stacked NES scores to permit comparison. Arrow heads indicate species-specific pathways. (8) Overlapping positive (left panel) and negative (right panel) gene set enrichments summarized in Venn diagrams. (C) Heatmap showing log_2_ expression of a subset of genes from the *IFNy response* gene set. Gene sets deemed to be significantly enriched if FDR-q < 0.05.

As for GSEA, a handful of *T. gondii* species-specific pathways identified were *Type II diabetes mellitus signaling* and *CD27 signaling in lymphocytes* and *H. hammondi*-specific immune-related pathways such as *GM-CSF* and *TGF-β signaling pathways* (Fig. 3Sa, Table S4).

### *T. gondii* and *H. hammondi*-infected cells have transcriptional profiles indicating distinct cell cycle states

Only two gene sets, *Myc targets v1* and *v2*, were significantly enriched in all *T. gondii*-infected cells regardless of strain type and either negatively enriched or not significantly enriched *H. hammondi*-infected cells (Figs. 3A and B, 3Sc (bottom panels) and 4A). Given the fact that *T. gondii* is known to induce *c-Myc* translocation into the nucleus and also induces S phase transition and ultimately G_2_/M cell cycle arrest in human host cells [37,46], we wanted to determine if the lack of induction in the *Myc targets v1* and *v2* gene sets in *H. hammondi*-infected cells reflected an inability of this species to manipulate cell cycle gene expression and ultimately progression through the cell cycle. We therefore looked specifically at transcript levels for a number of cell cycle related genes in *T. gondii*- and *H. hammondi*-infected cells and found multiple E2F-family related genes were of lower abundance in *H. hammondi*-infected cells compared to *T. gondii*-infected cells (e.g., *E2F1, E2F2* and *MCMs 4, 5, 6* and *7*; Fig. 4B and Table S3). This observation is consistent with the lower absolute NES values (for negative enrichment) in *H. hammondi* infection compared to *T. gondii* strains for the *G2M Checkpoint* and *E2F Targets* gene sets (Fig. 3A, Table S2), and suggests that *H. hammondi*-infected cells may be in a different cell cycle state. Consistent with this idea, we found that that *H. hammondi*-infected cells had significantly higher transcript abundance for *GADD45A*, *B* and *G* (genes involved in DNA damage-induced growth arrest) and *CDKN1A* and *CDKN1C* compared to *T. gondii*-infected cells (Fig. 4B), suggesting that *H. hammondi*-infected cells may have a phenotype consistent with DNA damage-induced cell cycle arrest. A role for activation of DNA-damage-mediated pathways is also supported by IPA, in which we identified 38 *H. hammondi*-specific pathways including *DNA damage-induced 14-3-3σ signaling* which contains a number of DNA damage-induced cell cycle checkpoint proteins [47,48] (Fig. 3Sb and Table S4; highlighted in blue). Combined with the differential *MYC targets* gene sets that were suppressed during *H. hammondi* infection compared to *T. gondii* (Figs. 3A and 4A), these data suggest that *T. gondii* and *H. hammondi*-infections may have a divergent impact on the cell cycle of the host cells that they infect. While *T. gondii* induces progression through S phase and into G_2_/M, *H. hammondi* infection may lead to cell cycle arrest (in either G_1_/S or G_2_/M) via DNA damage-related stress responses [49].

**Fig. 4.**
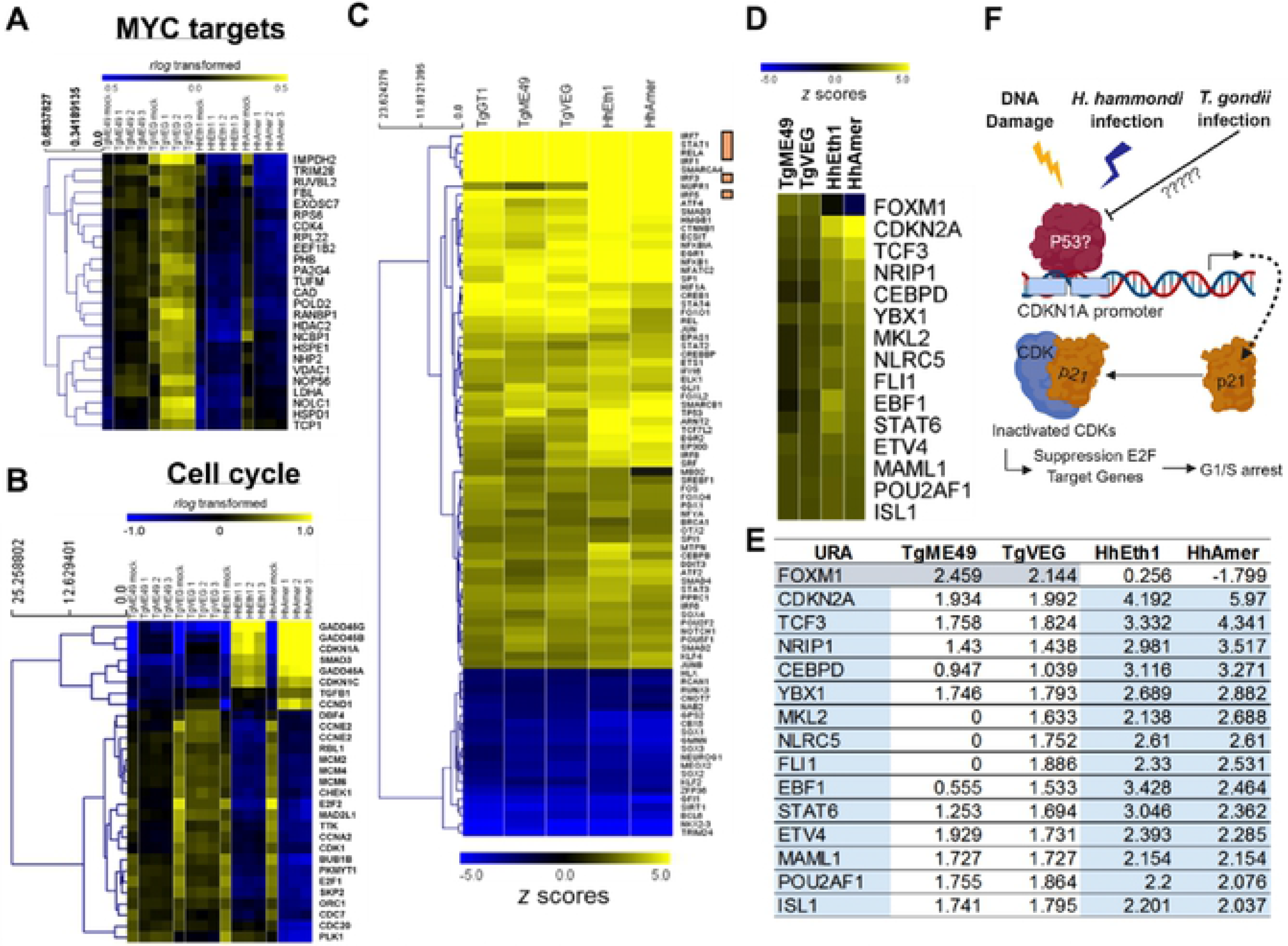
*H. hammondi* infection of THP-1 cells leads to a dramatically different impact on cell cycle regulation pathways. (A and B) Heatmaps showing log_2_ expression of a subset of genes positively enriched or depleted by T gondii (TgME49 and TgVEG) and H. hammondi (HhEth1 and HhAmer) in the MYC targets v1 (A) and cell-cycle gene sets (B; FDR-q < 0.05). Log_2_ normalized data were mean-centered amongst the samples and hierarchically clustered (Euclidean distance). (C) Heatmap of the z scores of common upstream transcription regulators significantly activated (yellow hues; z scores ≥ 2, *P*_*adj*_ < 0.01) or inhibited (blue hues; z scores s −2, *P*_*adj*_ < 0.01) in response to T. gondii (Tg) or H. hammondi (Hh) infections in THP-1 cells. Orange boxes highlight known T gondii altered host transcription factors. Heatmap (D) and table (E) showing z scores of upstream transcription regulators predicted to be activated (z scores ≥ 2) and/or inhibited (z scores ≤ −2) in parasites (T. gondii - grey; H. hammondi - light blue). Regulators were hierarchically clustered (Euclidean distances). (F) Schematic summary of the potential downstream impact of activation of the cyclin- dependent kinase (CDK) inhibitors CDKN1A and CDKN2A by H. hammondi based on pathway and upstream regulator analys is of RNAseq data (CDKN1A pathway shown). In response to H. hammondi infection, CDKN1A gene transcription increases and encodes p21. p21 can then bind to, and inactivate, CDKs, leading to reduced transcription of E2F target genes and cell cycle arrest. The most likely host initiator of this process would be P53, although other mechanisms of activation of this response are possible.

### Upstream regulator analysis reveals differential regulation of cell cycle-related transcription factors by *H. hammondi* infections

Using IPA upstream regulator analysis, we identified cascades of upstream transcription regulators that might be responsible for driving the observed transcriptional changes after infection with *T. gondii* and *H. hammondi*. We shortlisted upstream transcription regulators in response to *H. hammondi* or *T. gondii* infection (compared to mock-infected THP-1 cells) using *z* scores ≥ 2 for activation and ≤ −2 for inhibition (Fisher’s exact test; overlap *p* <0.01). From the analysis we identified type/strain-specific and common transcription regulators activated or inhibited in *T. gondii* and/or *H. hammondi* infections (Figs. 4C-E, Table S5). Common upstream transcription regulators include many well-known signaling mediators involved in *T. gondii* infection (e.g. multiple IRFs and STATs including IRF1, 3 and 5 and STAT1) as well as the NFκB p65-encoding gene *RELA* (Fig. 4C).

While most upstream regulators were shared between *T. gondii* and *H. hammondi*-infected cells (which was consistent with the number of infection-altered pathways shared between them at the transcriptional level), we did identify *FOXM1* as a significant *T. gondii*-specific upstream transcription regulator (average Z-score in *T. gondii* of 2.31 compared to −0.79 in *H. hammondi*), and *CDKN2A* as being much more highly enriched in *H. hammondi*-infected cells (average Z-score in *T. gondii* of 1.96 compared to 5.08 in *H. hammondi*; Figs. 4D and E). *FOXM1* is a component of the of the FOXM1-MMB complex which is active during G_2_/M and promotes transcription of E2F target genes, while *CDKN2A* (and *CDKN1A* as described above) are both involved in DNA-damage or stress-induced cell cycle arrest via CDK inhibition [50–53]. This provides further clues that *H. hammondi* might be regulating cell cycle progression differently than *T. gondii* during infection, and could be doing this in manner very similar to P53 mediated cell cycle arrest (Fig. 4F).

### *H. hammondi* and induced cytokine secretion *T. gondii*-infected THP-1 cells are in distinct phases of the cell cycle

We analyzed cell cycle progression of sporozoite-infected cells using flow cytometry. Consistent with the literature [37,46], a TgVEG-infected THP-1 cells showed a distinct cell cycle profile compared to uninfected cells, with a small but evident subpopulation of cells that were in G_2_/M (Fig. 5A). In contrast, HhAmer-infected cells looked much more similar to uninfected THP-1 cells with respect to the cell cycle (Fig. 5A). In a separate experiment we compared RH88-(type I *T. gondii* tachyzoites) infected THP-1 cells to THP-1 cells infected with HhAmer sporozoites, and again found that in contrast to *T. gondii*-infected THP-1 cells which exhibited a prominent G_2_/M peak, HhAmer-infected cells had a host cell cycle profile that lacked this peak and was much more similar to uninfected cells (Fig. 5B). All supernatants were assayed for CXCL10 secretion to verify *H. hammondi* infection (Fig. 5SA).

**Fig. 5.**
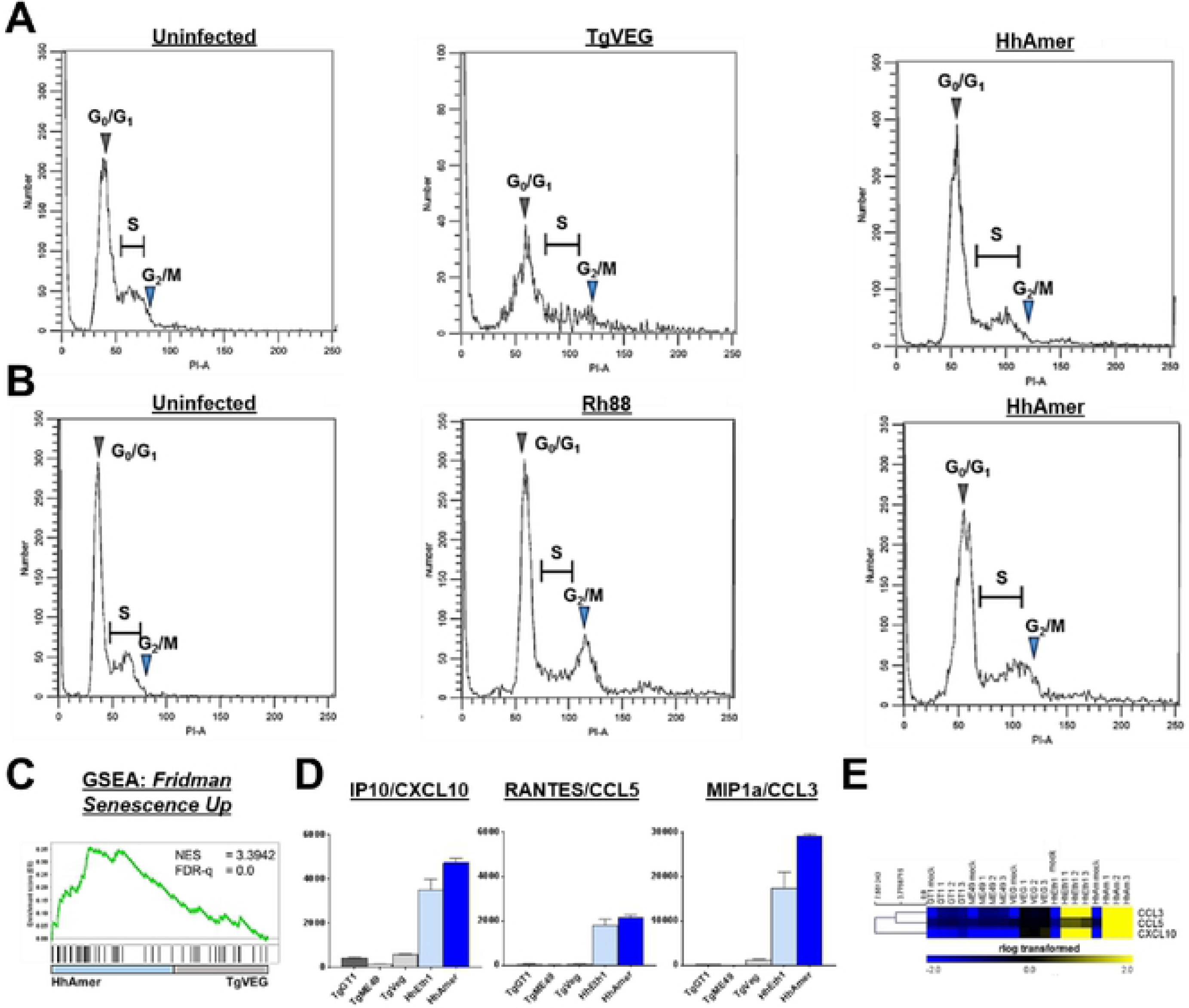
*H. hammondi-infected* THP-1 cells are arrested at the G_1_/S phase and displayed senescence-associated secretory phenotype. (A) THP-1 cells were infected with *T gondii* (TgVEG), *H. hammondi* (HhAmer) or uninfected for 20 hours and fixed with 80% ethanol. THP-1 cells were then stained with propidium iodide (Pl) and DNA content was analyzed with flow cytometry. Cell cycle progression was analyzed with ModFitLT software. Black lines represent histograms of DNA content (Pl-Area) vs. cell counts. Grey arrows show the G_0_/G_1_ peak and the G_2_/M peak (blue arrows) was determined at a ratio of 2 of the G_0_/G_1_ peak. Areas of histogram in between the G_0_/G_1_, and the G_2_/M peaks show host cells in the S phase. TgVEG-infected THP-1 cells displayed higher portions of cells found at the G_2_/M phase as compared to HhAmer- infected cells. (B) THP-1 cells were infected with *T gondii* (Rh88), *H. hammondi* (HhAmer) or uninfected for 20 hours and fixed with 80% ethanol. Host cells were stained as mentioned in A). Rh88-infected THP-1 cells displayed higher portions of cells found at the G_2_/M phase as compared to HhAmer-infected cells. (C) GSEA of the *Fridman Senescence Up* gene set in *T. gondii* (TgVEG-) and *H. hammondi* (HhAmer)- infected THP-1 cells. HhAmer-infected cells were enriched in the *Fridman Senescence Up* gene set with a normalized enrichment score (NES) of 3.3942. (D) Supernatants from THP-1 cells infected with *T gondii* or *H. hammondi* were collected 24 h post- infection and analyzed for pro-inflammatory cytokine secretion using the human cytokine 30-plex Luminex kit. Bars show relative changes in fluorescence intensity (fl) in parasite infected cells compared to mock-infected cells (fl infected - fl mock). Pro- inflammatory cytokines that were associated with SASP significantly different between *T. gondii* and *H. hammondi-infected* THP-1 cells are shown (One-way ANOVA Tukey's multiple comparisons test, *p*<0.05). Error bars represent SEM.(E) Heatmap shows mean-centered hierarchically clustered (Euclidean distance) rlog-transformed values of the respective cytokines shown in (D).

Among the gene sets that were significantly enriched in *H. hammondi*-infected THP-1 cells compared to *T. gondii* was the *Friedman Senescence Up* gene set [54] with an NES of 3.3942 (Fig. 5C), suggesting that *H. hammondi*-infected cells may have some similarities to senescent cells which would not only explain differences in the host cell cycle but also the increased production of inflammatory cytokines via the Senescence-Associated Secretory Phenotype (SASP) [55]. This hypothesis was further supported by IPA^®^ analyses showing significant enrichment of genes in the *DNA damage-14-3-3σ signaling* pathway (Fig. 3Sa), the higher expression of *CDKN1A*, increased *CDKN2A* signaling along with reduced *FOXM1* signaling (Fig 4). One of many defining characteristics of senescent cells is the production of lysosomal senescence-associated β-galactosidase (gal) [56–62] and β-gal enzyme activity can be detected both in cells and supernatants [63]. We evaluated this in parasite-infected THP-1 cells and found no significant increases in β-gal activity in THP-1 cells infected with either parasite species, even though β-gal activity did increase in Phleomycin-treated THP-1 cells (Fig. 5SB). However, we did find that supernatants from *H. hammondi*-infected THP-1 cells contained significantly higher levels of IP10/CXCL10 [64,65], RANTES/CCL5 [65–67], and MIP-1a/CCL3 [68,69], which are also well known SASP factors (Tukey’s multiple comparisons test \**p* <0.05), and this correlated with a similarly robust increase in transcript abundance for these chemokines (Figs. 5D-E and Table S6).

Using CXCL10 secretion as a sentinel chemokine for SASP, as expected we found that *H. hammondi* (HhAmer)-infected cells produced significantly more CXCL10 compared to mock-infected cells (2600 pg/10^5^ cells; Fig. 6A, left; Tukey’s multiple comparisons test, \*\*\*\**p*<0.0001), while supernatants from TgVEG-infected cells did not contain significantly higher CXCL10 compared to mock-infected cells (Fig. 6A; left). When Transwell^®^ inserts were used to separate parasites from THP-1 cells, CXCL10 induction by *H. hammondi* was completely abrogated (Fig. 6A), and heat-killed parasites of both species also failed to induce CXCL10 (Fig. 6B).

**Fig. 6.**
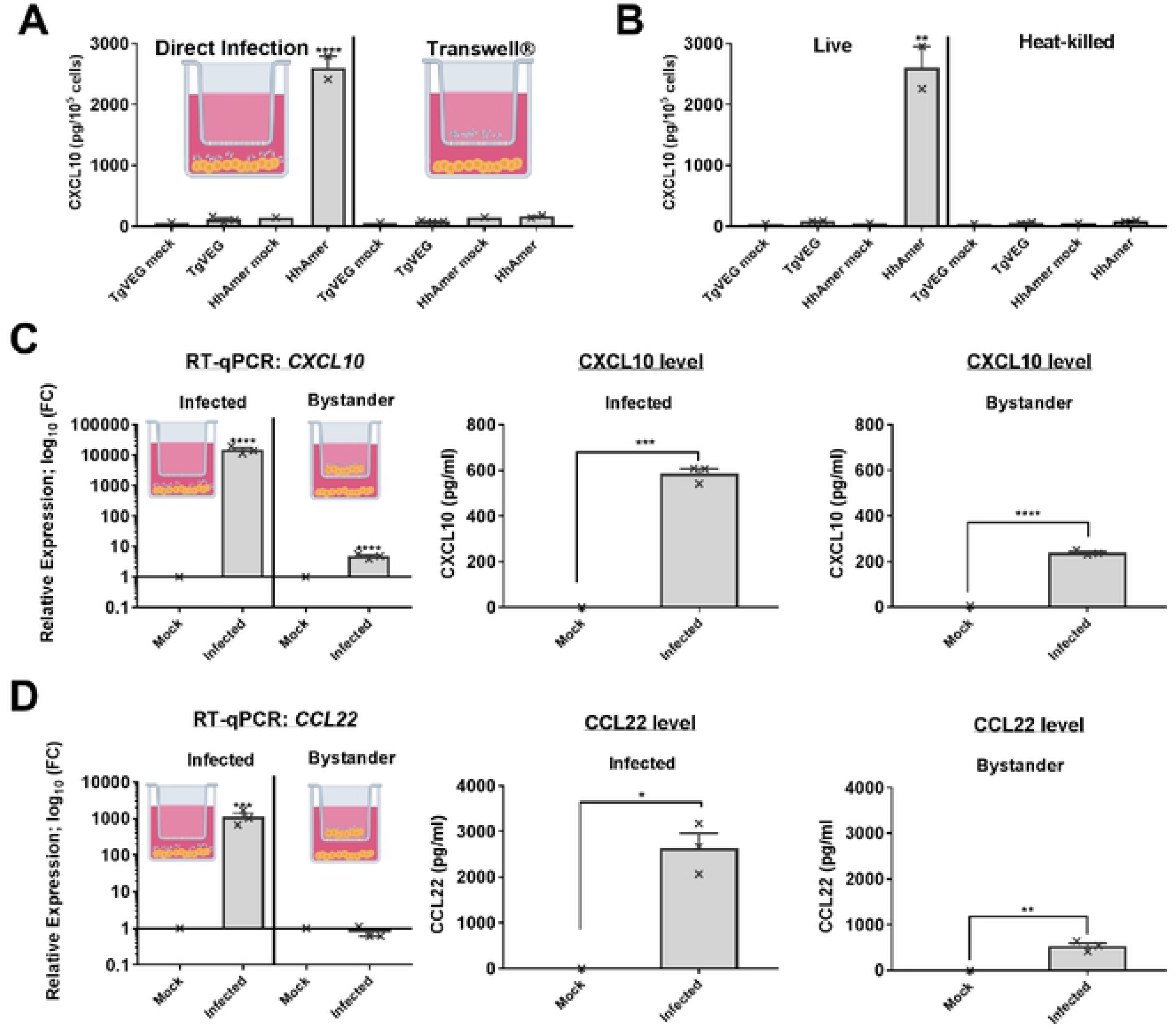
*H. hammondi* induction of CXCL10 requires active invasion and live parasites and possesses a paracrine effect. (A) *T. gondii* (TgVEG) or *H. hammondi* (HhAmer) sporozoites were added either directly to THP-1 cells or to Transwell^®^ inserts that separated the parasites from the cells. Secretion of CXCL10 was significantly different in HhAmer-infected THP-1 cells as compared to mock-infection control (****p<0.0001, Tukey's multiple comparisons test). (8) TgVEG or HhAmer were heat-killed at 95°C for 5 min and cooled to room temperature before exposing them to THP-1 cells. Secretions of CXCL10 were significantly higher in HhAmer-infected THP-1 cells as compared to mock infection (**p<0.01, Tukey's multiple comparisons test). THP-1 cells were infected with freshly excysted *T. gondii* or *H. hammondi* sporozoites at MOI of 2. Mock infection was performed by inoculating THP-1 cells with parasite preparation passed through a 0.2 µm filter. Supernatants from infected THP-1 cells were collected 24 h post-infection and analyzed for human CXCL10 secretion by ELISA. Error bars represent SEM. (C and D) THP-1 cells were seeded in the Transwell^®^ inserts and bottom of the wells. Freshly excysted *H. hammondi* Amer were added onto the Transwell^®^ seeded with THP-1 cells at MOIof 2. Supernatant and mRNA were collected from the bottom wells 24 h post-infection. Transcripts *(CXCL10* and *CCL22)* and protein (CXCL10 and CCL22) levels were analyzed with RT-qPCR and ELISA respectively. The phenotype of *H. hammondi-mediated* chemokine induction during live parasite invasion was confirmed (infected). Direct *H. hammondi* infection induces significant ly higher chemokine transcripts and protein levels in THP-1 cells (Tukey's multiple comparisons test, \**p*<0.01,\*\*\**p*<0.001 and ****p<0.0001).Transcript and protein levels of CXCL10 were significantly higher in both the bystander cells in the bottom wells not exposed to direct parasite infection as compared to their respective mock-infected THP-1 cells (unpaired *t* test, \*\**p*<0.01,and ****p<0.0001). Unlike THP-1 cells exposed to direct parasite infection, transcript of *CCL22* was not significantly higher in cells in the bystander cells not exposed to direct parasite infection (Tukey's multiple comparisons test, ***p<0.001). The level of CCL22 secretions were significantly different in the infected and bystander cells as compared to their respective mock-infected THP-1 cells (unpaired *t* test, \**p*<0.05 and \*\**p*<0.01). Bar graphs represent mean of cytokine secretions and crosses represent each individual data points.

The SASP can be transferred to bystander cells by paracrine signaling [70], and to test this we used Transwell^®^ inserts to separate *H. hammondi*-infected THP-1 cells from bystander and quantified CXCL10 induction by qPCR. We found that bystander cells exposed to HhAmer and host excretory/secretory products (ESPs) from the Transwell^®^ insert expressed significantly higher *CXCL10* transcript as compared to bystander cells of the Transwell^®^ insert of the mock-infected THP-1 cells (by ~10-fold; Fig. 6C; Tukey’s multiple comparisons test, \**p*<0.01, \*\*\**p*<0.001 and \*\*\*\**p*<0.0001). RT-qPCR for *H. hammondi GRA1* transcript confirmed the absence of parasites in the bystander cells (Table S7), and shows that CXCL10 transcript induction was due exclusively to a paracrine effect from infected THP-1 cells from the upper chamber of the Transwell^®^ insert. CXCL10 was detected in supernatants in both conditions, since this chemokine could freely diffuse after its secretion. In contrast to CXCL10, we did not observe any increase in transcript abundance for *CCL22* in bystander cells (Fig. 6D). CCL22 is also induced by *T. gondii* and *H. hammondi* infection [24], but is not a typical constituent of the SASP [71]. Taken together this provides further evidence for as SASP-like response in *H. hammondi*-infected, but not *T. gondii*-infected, THP-1 cells.

### *T. gondii*- conditioned medium suppresses *H. hammondi*-induced cytokine secretion in THP-1 cells and increases replication rates of *H. hammondi* in primary human fibroblasts

Next, we speculated that *T. gondii* might be capable of suppressing the secretory response by *H. hammondi*. We exposed THP-1 cells to TgVEG sporozoites, or to 4 h conditioned medium from TgVEG-infected THP-1 cells, for 4 h prior to infection with *H. hammondi*. Interestingly we found that TgVEG sporozoites and TgVEG-conditioned media both suppress *H. hammondi*-mediated cytokine secretion (Figs. 7A and B). However, when THP-1 cells were pre-infected with *H. hammondi* sporozoites, *T. gondii*-mediated cytokine suppressive effect was no longer observed (Fig. 7A).

**Fig. 7.**
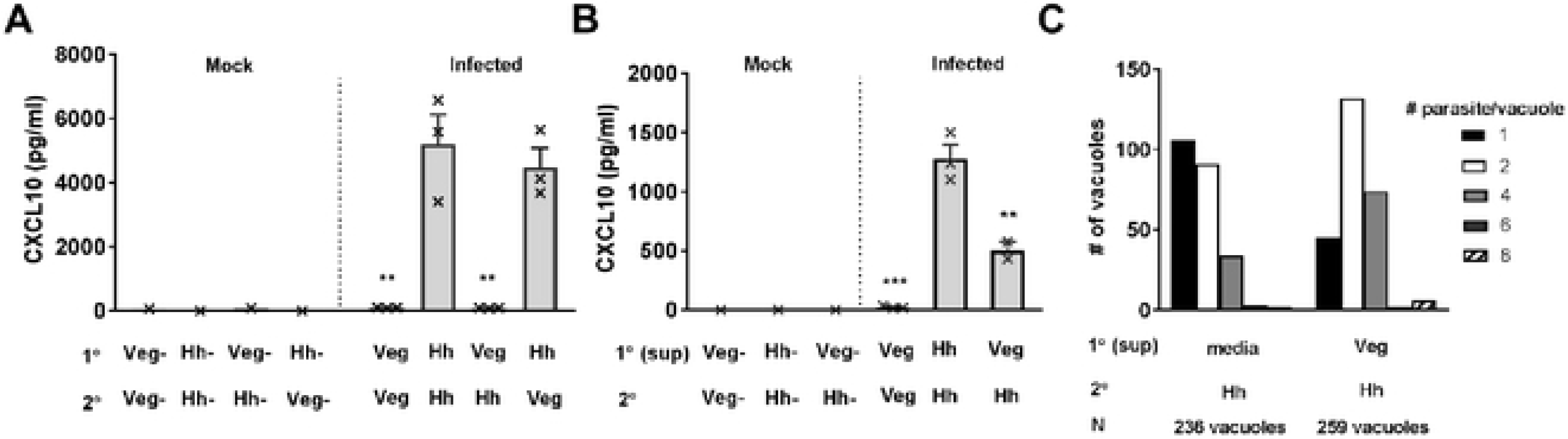
*T. gondii-mediated* suppression of chemokine increases replication rates of *H. hammondi.* (A) THP-1 cells were pre-infected (1°) with an MOI of 2 of freshly excysted *T. gondii* (Veg) or *H. hammondi* (Hh) sporozoites. 4 h post-pre-infection, an MOI of 2 of freshly excysted Veg or Hh were added into the same wells (2°). Supernatant was collected at 20 h post 2° infection and CXCL10 level was analyzed with ELISA. Bar graphs show mean concentration of CXCL10 (pg/ml) and crosses represent each individual data points of the biological replications. CXCL10 secretion of THP-1 cells pre-infected with Veg was significantly lower than THP-1 cells pre-infected with Hh regardless of the species of parasites for the 2° infection (Sidak’s multiple comparisons test, \*\**p*<0.01 compared against Hh (1°)-Hh (2°). (B) Primary infection extract (1° (sup)) was prepared by infecting THP-1 cells with an MOI of 2 of freshly excysted Veg or Hh sporozoites for 4 h. Supernatant was collected from the cells and filtered through a 0.2 µm filter (ESP). New wells of THP-1 cells were pre-infected with the ESP for 4 hr (1° (sup)). Next, an MOI of 2 of freshly excysted parasites were added into the same wells (2°). Supernatant of the cells were collected 20 h post 2° infection and CXCL10 level was analyzed with ELISA. Secretion of CXCL10 was significantly lower in THP-1 cells pre-infected with Veg ESP regardless of the parasite species for the 2° infections (\*\**p*<0.01, \**p*<0.001; Tukey's multiple comparisons test against Hh (1° sup)-Hh (2°)). (C) Histogram showing vacuole size of *H. hammondi* growth in the absence and presence of *T. gondii* (VEG) and THP-1 cells ESP.Total numbers of vacuoles (N) were counted from two coverslips at 72 h post 2° infection.Veg and THP-1 cells ESP was prepared in THP-1 cell infections as described in (B). HFF cells were pre-conditioned with Veg and THP-1 cells ESP and freshly excysted Hh sporozoites were added into the same wells. Growth rates of parasite on HFF cells pre-conditioned with ESP was significantly higher than HFF cells that did not pre-condition with ESP (\**p*<0.05, unpaired *t* test comparing means of vacuole size of the coverslips; means of unconditioned vs. conditioned well are 2.141±0.1479 and 2.869±0.2169 respectively).

Our previous work found that *H. hammondi* replicates ~4-8-fold slower than *T. gondii* [34], and given the divergent effects on the host cell cycle we hypothesized that *T. gondii* infection may result in a more permissive host cell state, and that this may alter the replication rate of *H. hammondi*. To address this we infected HFF cells that were pre-treated with TgVEG/THP-1 conditioned medium (collected 4 h post THP-1 infection) with freshly excysted HhAmer sporozoites and quantified parasite vacuole number and size. Remarkably, we observed a significant increase in *H. hammondi* vacuole size in pre-conditioned HFFs compared to untreated HFFs in two separate experiments performed with completely distinct parasite batches and reagents (Figs. 7C and 7S; \**p*<0.05, unpaired *t* test comparing means of vacuole size of the coverslips; means of unconditioned vs. conditioned well are 2.141±0.1479 and 2.869±0.2169 respectively). In particular we saw a decrease of vacuoles with single parasite and increase of two, four and eight parasites per vacuole in the TgVEG/THP-1 conditioned medium (Fig. 7C). While there was variation in the impact of TgVEG-conditioned media on *H. hammondi* replication (data in Fig. 7S show a much more dramatic increase in vacuole size compared to data in Fig 7C), these data clearly show that *T. gondi*-infected host cells produce (a) factor(s) that can render bystander cells more permissive to parasite replication (at least for *H. hammondi*), and that the replication rate of *H. hammondi* can at least be partially determined by the condition of the host cell.

## Discussion

It is well known that *T. gondii* tachyzoites activate a potent host immune response. Despite that, once inside a host cell, *T. gondii* is able to modulate, survive and even evade immune responses, all of which lead to replication and ultimate dissemination to distant host tissues. This has largely been attributed to the powerful strategy that *T. gondii* employs to co-opt host gene expression and to protect itself from host proteins designed to destroy these parasite-containing vacuole [15–17,72,73]. In contrast little is known about the impact of *H. hammondi* on the host cell, and whether its comparatively “avirulent” lifestyle can be attributed at all to differences in the host response. Many of the unique aspects of *H. hammondi* biology, including its comparatively slower replication rate, spontaneous (and terminal) cyst formation and inability to be lethal even in IFNγ knockout mice [29,34,74], suggest that it is engaged in an inflexible developmental program once sporozoites are released from the oocyst. In the extreme, then, one could hypothesize that the nature and magnitude of the host response may be completely irrelevant to the outcome of infection with *H. hammondi*. However our data provide strong evidence to the contrary.

At first glance, our comparisons of the transcriptomes *T. gondii* and *H. hammondi*-infected host cells indicate that these parasites modulate the host cell using qualitatively similar strategies. Over 95% of the underlying altered pathways were shared between *T. gondii* and *H. hammondi*, and fell into categories like inflammation, apoptosis, cell growth, and metabolism that have long been known to be targeted by *T. gondii* [4–6] (Fig. 3). However when the magnitude of the response was taken into account (whether measured by normalized enrichment scores from GSEA or cytokine secretion), it is clear that *H. hammondi*-infected cells are responding much more robustly to infection compared to those infected with *T. gondii*. This effect, which is also recapitulated in an animal model of infection when differences in parasite burden is taken into account (Fig. 2), suggests either that a) *H. hammondi* has fewer countermeasures to counteract the host response than *T. gondii* or b) *H. hammondi* has a factor or factors that cause hyperactivation of the immune response (via secreted effectors, and/or production of pathogen- or damage-associated molecular patterns).

Intriguingly our results show that the most obvious differential host response in THP-1 cells infected with *T. gondii* and *H. hammondi* is the regulation of cell cycle (Figs. 3–5), a phenomenon long-described in *T. gondii* but whose role in determining infection outcome is unknown. Unlike *T. gondii* infection that actively induces host *c-Myc* [75], the *MYC targets v1* and *v2* gene sets were suppressed in THP-1 cells infected with *H. hammondi* (Figs. 3 and 4). *T. gondii* infection leads to an accumulation of cells in the G_2_/M phase [37,46] and in our data this was accompanied by increased transcript abundance of *FOXM1*-transcribed genes (Fig. 4; [50,52]). In contrast, *H. hammondi* leads to increased transcription of multiple *GADD45* genes which block cell cycle progression as a part of the DNA damage response [76–78]. DNA damage can induce P53 activation and increase transcription of *CDKN1A*, which we observed only in *H. hammondi*-infected cells (Fig. 4D-F; [48]). DNA damage responses can have dramatic effects on cell biology that are independent of cell cycle arrest, and one of these is the Senescence Associated Secretory Phenotype (SASP; [65,70,71]). We observed increased production of multiple chemokines typically associated with the SASP, including CXCL10 which not only was uniquely present in *H. hammondi*-infected supernatants but was also activated in bystander cells via paracrine signaling (Figs. 5 and 6). We did not detect elevated β-gal activity in host cells infected with *H. hammondi,* cell types and different cellular conditions could also affect the reliability of any types of senescence markers [79]. It is very intriguing to postulate that the manipulation of the host cell cycle by *T. gondii*, and specifically its ability to induce S-phase transition and ultimately G_2_/M arrest, is a means to prevent the host cell from becoming senescent and activating the SASP in both the infected cell and ultimately in neighboring bystander cells. Disrupting the ability of the host cell to secrete cytokines like CXCL10 would certainly be advantageous given the importance of this chemokine to maintain T-cell populations capable of controlling *T. gondii* proliferation [41,80].

Similar to paracrine effects of the *H. hammondi*-induced SASP on bystander cells, *T. gondii* induces host- and infection-altering factors to be produced by host cells as well. *H. hammondi* has a markedly slow replication rate compared to *T. gondii* (shown in multiple studies including [34,74]), but this slow rate of division appears to be flexible depending upon the host cell environment. The ability to alter the replication of an intracellular parasite like *H. hammondi* using *T. gondii*-conditioned medium (Fig. 7) leads to a number of interesting biological questions about the nature and identity of the factors capable of controlling parasite replication. Supernatants from *T. gondii*-infected cells have been found to have a variety of effects on recipient cells that vary depending on the donor and recipient cell types (e.g., [81,82]). Although to date the factors themselves have not been identified, interaction between *T. gondii* and the host cells which reduced UHFR1 and cyclin B1 has been shown to be advantageous for *T. gondii* replication [46]. Our data provide further compelling evidence for their importance in determining the outcome of *T. gondii* infections. In addition to this interesting biology there are additional practical implications of this work with respect to the *H. hammondi*/*T. gondii* comparative system. The use of *T. gondii*-conditioned media may be a first step in further improving culture conditions for *H. hammondi*, with the ultimate goal of promoting long term *in vitro* culture which is not yet possible [29,74].

## Materials and Methods

### Cells

Human foreskin fibroblasts(HFFs) and human monocytes (THP-1) were maintained in cDMEM (100 U/ml penicillin/streptomycin, 100 μg/ml streptomycin, 2 mM L-glutamine, 10 % FBS, 3.7 g NaH_2_CO_3_/L, pH7.2; ThermoFisher Scientific) and cRPMI-1640 (100 U/ml penicillin/streptomycin and 10 % FBS; ThermoFisher Scientific) respectively. All cells were grown at 37 °C in 5 % CO_2_. One day prior to parasite infection, THP-1 cells growth media were replenished with 20 % (v/v) of new media (cRPMI). All THP-1 cell infections were performed in cDMEM.

### Mice

Balb/c mice were purchase from Jackson Laboratory and were female aged 6-8 weeks. All animal experiments were approved by the local IACUC at the University of Pittsburgh (Protocol #18032113 and #092011), with euthanasia and anesthesia conducted according to AVMA guidelines. Anesthesia of the rodents were performed with CO_2_ followed by decapitation of the animals. Euthanasia was performed with isofluorine.

### Parasites

Oocysts of *Toxoplasma gondii* (Tg) genotype I (GT1), II (ME49) and III (VEG) and *Hammondia hammondi* (Hh) American (Amer) and Ethiopian-1 (Eth1) isolates were harvested from cat feces 7-11 days after feeding mouse tissues (brain for *T. gondii*, leg muscle for *H. hammondi*) infected with parasites to 10-20 week old cat free of pathogens [83,84]. Unsporulated oocysts were isolated via sucrose floatation and placed at 4 °C in 2 % H_2_SO_4_ to encourage sporulation and for long-term storage.

### *T. gondii* and *H. hammondi* oocyst excystation

Sporulated oocysts were washed 3 X in Hank’s balanced salt solution (HBSS; Life Technologies) and treated with 10% bleach (in PBS) for 30 min with shaking at room temperature. Next, bleach was washed off using HBSS and parasites were added to 4 g sterile glass beads (180 μM; Sigma-Aldrich). Oocysts were vortexed on high speed for 15 s on/15 s off for a total duration of 2 min to disrupt the oocyst wall mechanically. Sporocysts were pelleted by centrifugation at 1,000 x g for 10 min. Pellet was resuspended in 5 ml of pre-warmed and freshly made, 0.22 μM filtered excystation media (0.1 g porcine trypsin (Sigma-Aldrich), 2 g Taurocholic Acid (Sigma-Aldrich) in 40 ml PBS, pH 7.5). Sporocysts were incubated in 37 °C water bath with 5 % CO_2_ for 45 min and syringe-lysed using a 25 and a 22 gauge needle. *H. hammondi* sporocysts were syringe-lysed with a 25 gauge needle. To quench the excystation media, 7 ml of cDMEM was added. Excysted parasites (sporozoites) were pelleted and resuspended in cDMEM and grown onto monolayers of HFFs for overnight at 37 °C in 5 % CO_2_. Freshly excysted sporozoites were used in *in vivo* and experiments analyzing induction of CXCL10 in Transwell^®^, heat-killed assays and parasite growth assays.

### Human monocyte cell line infection

Prior to infection, THP-1 cells were seeded at 1 × 10^5^ cells/well in 24-well plates in cDMEM. To prepare parasites for the infections, HFFs monolayers containing parasite sporozoites were scraped, syringe-lysed, and pelleted (in some cases, freshly excysted oocysts were used). After resuspending pellet in cDMEM, the parasite mixture was filtered through a 5 μM syringe-driven filter (Millipore). THP-1 cells were infected with either *T. gondii* or *H. hammondi* at multiplicity of infection (MOI) of 4, 2 or 1.6. Three biological replicates were made for each parasite infection. THP-1 cells were also mock-infected with parasite filtered through a 0.22 μM syringe-driven filter (Millipore).

### RNA isolation

RNA was collected at 24 h post-infection from parasite- and mock-infected cells using the RNeasy Kit according to the manufacturer instructions (Qiagen). QIAShredder spin columns (Qiagen) were used to homogenize the samples prior to RNA extraction and contaminating DNA was degraded using RNase-free DNase (Qiagen). RNA was eluted in 50 μl of RNase-free water and gel electrophoresis and Nanadrop RNA quantification were performed to ensure RNA integrity. One mock and three biological infection replicates were done for all experiments. Total RNA samples were kept at −80 °C.

### mRNA-sequencing and data processing

mRNA-sequencing libraries and Illumina next generation sequencing were performed at the Core Facility at the University of Pittsburgh. Integrity of the RNA was analyzed with Agilent 2100 Bioananalyzer and all purified RNA samples had RIN scores >9. RNA was sequenced using NextSeq 550 (Illumina) and were pooled and sequenced over four lanes. Strand-specific, 150 bp, single-end RNA-sequencing was performed. Read libraries were mapped to the human genome (*Homo sapiens* ensemble v81; hg38) and transcriptome (*Homo sapiens* ensemble v81; hg38) with default options on CLC Genomics Workbench v11.0. Fastq files have been deposited in the NCBI short read archive (Accession number: SRX3734421-8, and accession numbers pending for some of the Fastq files).

### Differential expression analysis using *DESeq2* and Pre-ranked Gene Set Enrichment Analysis

*DESeq2* [40] was used to perform differential expression analysis of genes in THP-1 cells infected with *T. gondii* and *H. hammondi*. Raw reads (total gene reads; exported from CLC Genomics Workbench) with at least 1 read count in all samples were analyzed. Prior to analyzing differential gene expression, integrity of the data was examined using principal component analysis (PCA) and distances of all samples were calculated (embedded in the *DESeq2* package). Genes were considered to be differentially expressed in THP-1 cells if the log_2_ fold-change was ≥ 1 or ≤ −1 and with a *p*_*adj*_ value (alpha) <0.01.

Pre-ranked Gene Set Enrichment Analysis (GSEA; [45]) was performed to compare gene sets that were enriched in THP-1 cells in relation to parasite infections. Ranked list calculating the fold-change difference (subtracting normalized log_2_ fold-change of infected host cells from mock infected host cells) was used for the analysis.

### Ingenuity^®^ Core Pathway Analysis

Core analysis from Ingenuity^®^ Pathway Analysis (IPA; Qiagen) was used to examine biological relevance of the RNA-seq data. Canonical pathways and upstream regulators that are over-represented in *T. gondii* and *H. hammondi* infection were analyzed using log_2_ fold-change and *p*_*adj*_ value obtained in *DESeq2* [40]. Only genes that were deemed significantly expressed were used in the analysis (log_2_ fold-change ≥ 1 or ≤ −1 and *p*_*adj*_ value < 0.01 for Infected vs. Mock). IPA default settings were used for the analysis. For both canonical pathway and upstream regulator analysis, pathways and genes with activation *z* scores ≥ 2 were deemed to be activated while activation *z* scores ≤ −2 were deemed to be inhibited for different analysis. Threshold for *p* value significant used in the analysis was <0.05 or −log(p-value) >1.3.

### Mouse peritoneal cell collection

For mouse infections, 40,000 24 h parasite zoites or 20,000 freshly excysted sporozoites (TgVEG or HhAmer) in 200 μL of PBS were intraperitoneally injected into 7-9 week old mice. Mice were euthanized with CO_2_ according to IUCAC protocols and dissected. Peritoneal cells were collected in 3 ml PBS as described previously [85].

### Multiplex cytokine analysis

THP-1 cells were infected with MOI 4 of *T. gondii* or *H. hammondi* (same infection set up as the RNA-seq experiments). THP-1 cells infected or mock-infected with parasites were pelleted at 1,000 x g for 10 min. Supernatant were collected and stored at −80 °C. Luminex was performed at the Luminex Core Facility University of Pittsburgh Cancer Institute. Each sample was measured (fluorescence intensity) in duplicate.

### Cytokine analysis using ELISA

Concentrations of the human pro-inflammatory cytokine CXCL10 and CCL22 and mouse Ifnγ, Il12p40, Cxcl10 and Ccl22 in mouse peritoneal cells were analyzed by ELISA according to the manufacturers instructions (Human and Mouse DuoSet ELISA respectively, R&D Systems). For relative chemokine concentration in comparison to parasite load, log_2_ values of the absolute concentration of chemokines was calculated and relative expression in relation to parasite burden was calculated using the 2^−ΔΔCT^ method (ΔΔC_T_ = log_2_ chemokine concentration – ΔC_T_ *GRA1*).

### Transwell^®^ and heat-killed parasite infections

Freshly excysted TgVEG or HhAmer was added to THP-1 cells seeded in 24-well plates (control) or added onto Transwell^®^ inserts (Corning). For heat-killed parasite infection, freshly excysted sporozoites were heat-killed at 95 °C for 5 min and cooled to room temperature before infecting THP-1 cells. Multiplicity of infection of 2 was used in these experiments. Supernatants were collected either by pelleting the cells or collected from the bottom chamber of the Transwell^®^ setup.

### Reverse transcriptase-quantitative PCR

RNA were extracted using RNeasy RNA extraction kit as above and according to manufacturer’s instructions (Qiagen). cDNA was reverse transcribed from 1 μg of RNA using SuperScript IV First-Strand synthesis system (ThermoFisher Scientific). RT-qPCR was performed using QuantStudio with SYBR Green (Applied Biosystems). The PCR mixture contained 1 X SYBR Green buffer (BioRad), 0.25 μl of forward and reverse primers (Table S8) and cDNA. Genes were amplified using the standard protocol (95 °C for 10 min and 40 cycles of 95 °C for 15 sec and 60 °C for 1 min). Data were acquired and analyzed using QuantStudio™ Design & Analysis Software (ThermoFisher Scientific) and data exported to Microsoft Excel for threshold values (C_T_) and 2^−ΔΔCT^ for fold change analysis. GAPDH and *Gapdh* was used as the human and mouse reference gene respectively (control) for the 2^−ΔΔCT^ analysis. For *in vivo* 2^−ΔΔCT^ analysis, relative expression of target genes were normalized against ΔΔC_T_ of mock-infected mice. For relative expression normalized by comparison with parasite load, we used *GRA1* and as a reference gene for parasite burden. To do this, the ΔC_T_ of target genes and *GRA1* were first obtained (C_T_ target gene or *GRA1* – C_T_ *Gapdh*). Then, ΔΔC_T_ target gene was calculated (ΔC_T_ target genes – ΔC_T_ *GRA1*). Relative expression was finally calculated using the 2^-ΔΔCT^ methods for respective target genes. Primers validation and melt curve analysis were performed to ensure integrity of the RT-qPCR. No RT and water controls were also included in every RT-qPCR.

### Propidium iodide cell cycle analysis

Cell cycle profiles of THP-1 cells were analyzed with propidium iodide staining according to manufacturers instructions (ThermoFisher Scientific). THP-1 cells were infected with *T. gondii* or *H. hammondi* at an MOI of 2. For uninfected cells, THP-1 cells were seeded at the same time as the other cells in different treatment groups and an equal volume of media was added to the uninfected cells in place of parasite infection. At 20 h post-infection, cells were pelleted and washed 1 X with chilled PBS and fixed in chilled 80% ethanol at −20 °C for overnight. Cells were then washed with chilled PBS and rehydrated for 15 min with chilled PBS. Cells were stained with 2 μg/mL propidium iodide/RNaseA (ThermoFisher Scientific) solution for 15 min at room temperature with gentle agitation. Stained cells were analyzed immediately by flow cytometry using LSR II (BD Biosciences) at the United Flow Core at University of Pittsburgh or kept at −20 °C in the dark for no longer than 24 hr until flow cytometry analysis. Cell cycle profiles were analyzed with ModFit LT 5.0 (VSH).

### β-galactosidase activity assay

β-galactosidase activity was analyzed in THP-1 cells and secreted proteins using the β-galactosidase assay kit (ThermoFisher) according to manufacturers instructions. Briefly, THP-1 cells were inoculated with *T. gondii* or *H. hammondi*, treated with 1 μg/mL of phleomycin in cDMEM. At 72 h post-infection, cells were pellected at 300 x g for 10 min and supernatant was collected for β-galactosidase activity assay. The cell pellets were washed 2 times with PBS and lyzed with IP lysis buffer (ThermoFisher) with Halt Protease Inhibitor Cocktail (ThermoFisher), before analyzing for β-galactosidase activity. β-galactosidase activity was determined after 1 h of incubation at 37 °C and absorbance was read at 420nm.

### *H. hammondi* growth (vacuole size) under *T. gondii*/THP-1 conditioned medium

*T. gondii* Veg and THP-1 cells ESP was prepared in THP-1 cell infections as described above. Briefly THP-1 cells were infected with *T. gondii* VEG with an MOI of 2 for 4 h. HFF cells pre-seeded on coverslips were pre-conditioned with TgVeg/THP-1 cells ESP and freshly excysted Hh sporozoites were added into the same wells. At 72 h post-infection, coverslips were washed 2 times with PBS and fixed with 4% paraformaldehyde and stained with DAPI. The total number of visible vacuole and number of parasites per vacuole was counted from two coverslips for each infection conditions. Differences in the growth rates of the two conditions were determined by calculating the mean vacuole size of each coverslips.

## Acknowledgments

The authors thank Cori Richards-Zawacki and Carolyn Coyne for sharing cells and equipment.

---

**Fig. 2S.**
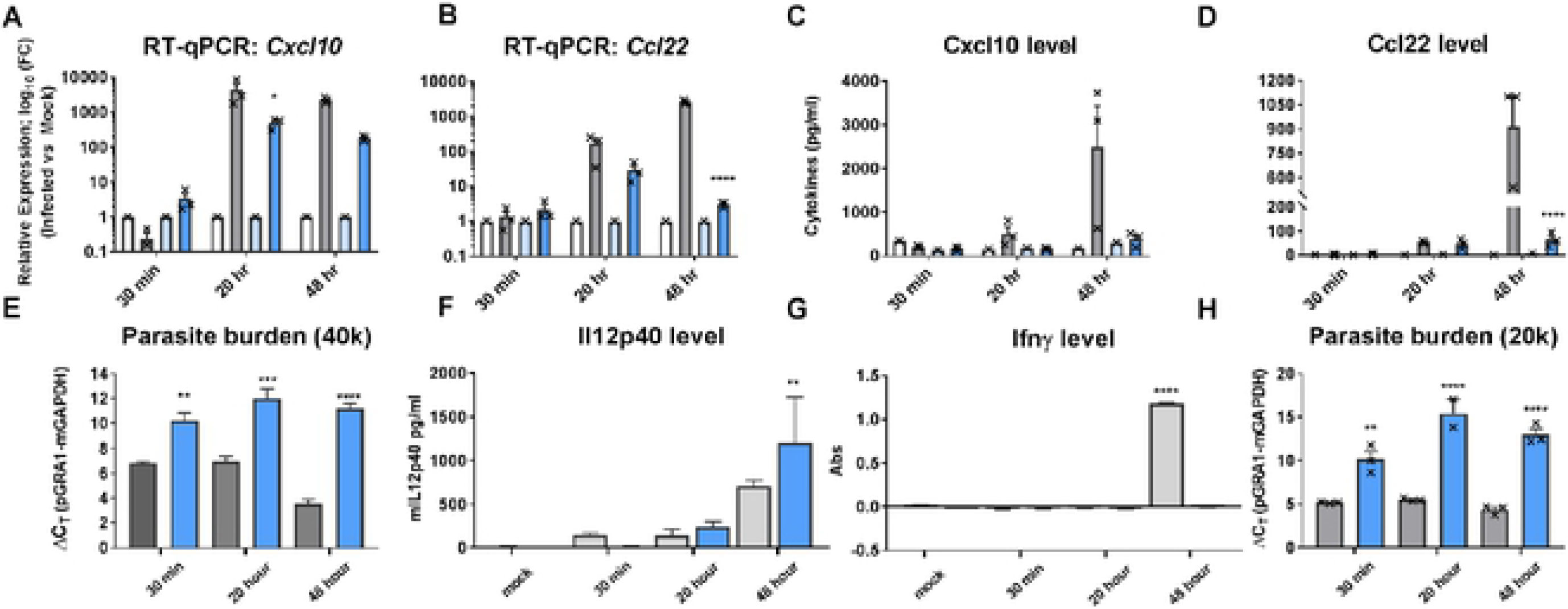
*T. gondii* VEG (TgVEG) and *H. hammondi* Amer (HhAmer) sporozoite infections *in vivo*. Mice were infected with T gondii (TgVEG; grey) or H. hammondi (HhAmer; blue). Mice were also mock-infected with parasite free extracts (TgVEG-free extract; white or HhAmer-free extract; light blue). Mouse peritoneal cell RNA and supernatants were collected at 30 min, 20 and 48 h post sporozoite infections. mRNA levels of chemokines (Cxcl10 and Ccl22) and parasite GRA 1 genes were quantified using RT-qPCR and Gapdh was used as the reference gene. Protein levels were quantified using ELISA. (A-8) Bar graphs show relative expression (RE) of chemokine genes *Cxcl10* (A) and *Ccl22* (B) calculated using the 2-MCT method (Infected vs. Mock). * represents significant RE of host responses in TgVEG as compared to HhAmer in the respective time points (Sidak's multiple comparisons test, \**p*<0.05 and ****p<0.0001). (C-D) Cxcl10 and Ccl22 protein levels were quantified and compared to mock-infected mice. Cxcl10 levels were not significantly different between species at all time points (Sidak's multiple comparisons test; *p*>0.05). Production of Ccl22 was significantly higher in TgVEG-infected mouse peritoneal cells at 48 h post-infection (Tukey's multiple comparisons test; ****p<0.0001). (E & H) Parasite loads in relative to *Gapdh* reference gene) in mice infected with 24 h post-excystation 40,000 (A) and 20,000 freshly excysted (D) TgVEG or HhAmer sporozoites. Bar graphs show LlCT (C_T_*GRA 1* - C_T_ *Gapdh)* at three different time point post-infections. Parasite burdens of HhAmer are lower than TgVEG (smaller LlCT values observed in mice infected with TgVEG; Sidak’s multiple comparisons test "*p<0.01, ***p<0.001 and ****p<0.0001 as compared to the respective time points). Secretions of 1112p40 (F) and lfny (G) in mouse infected with TgVEG or HhAmer. Mice were also mock-infected with parasite free extracts and samples collected at 48 h post-infection. Supernatants from mouse peritoneal cells were collected at the same time points as in (A). While secretions of 1112p40 were significantly higher in HhAmer-infected mice 48 h post-infection (Tukey’s multiple comparisons test ***p<0.01), secretions of 1112p40 were not significantly different in mice infected with TgVEG and HhAmer at any time points (Tukey's multiple comparisons test; *p*>0.05). Secretion of lfny was significantly higher in TgVEG 48 hour post-infection as compared to the other time points (Sidak's multiple comparisons test ****p<0.0001). Crosses show individual data point. Error bars show standard error of mean (SEM).

**Fig. 3Sa.**
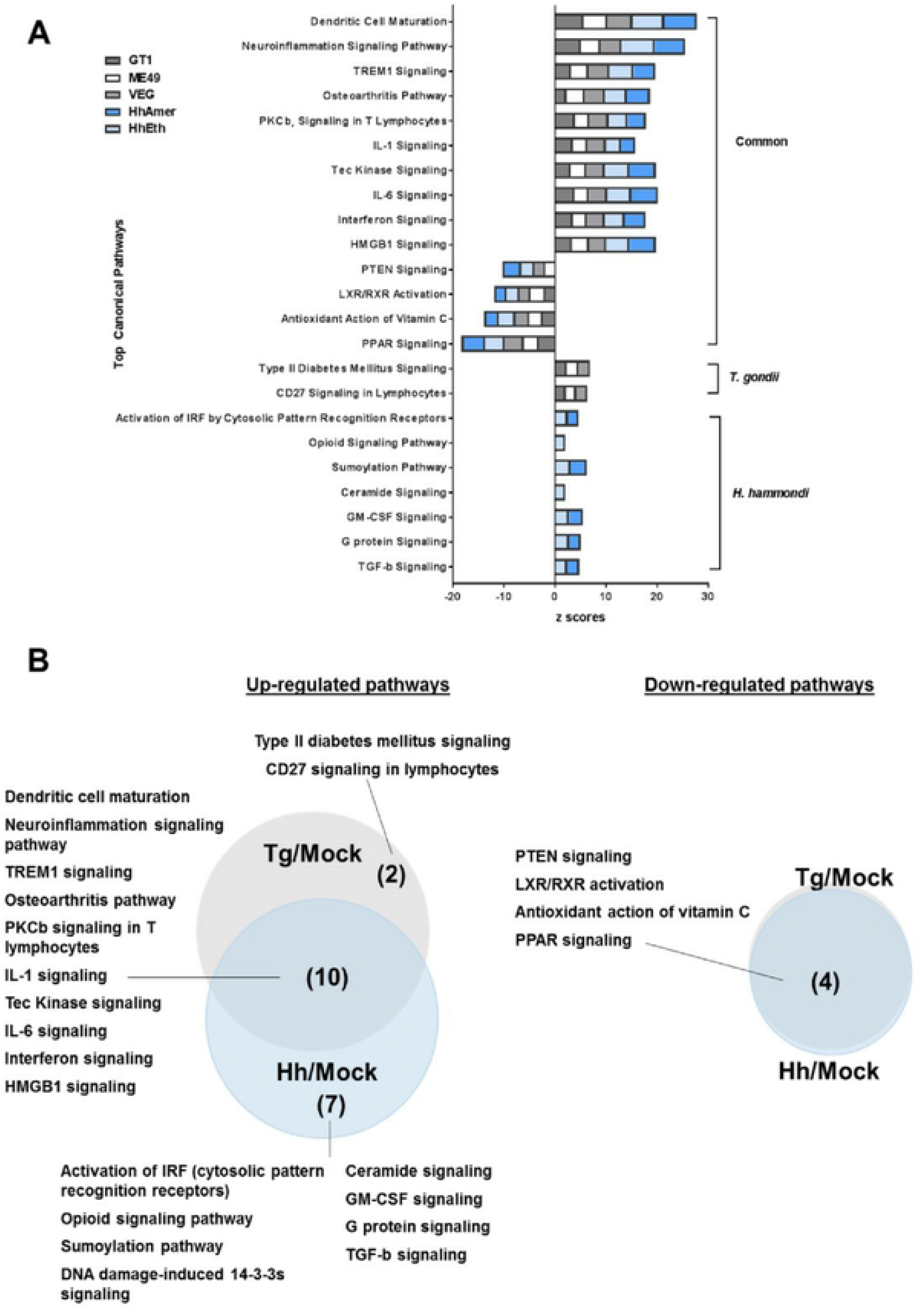
Top canonical pathways enriched (z scores) in *T. gondii* or *H. hammondi-infected* THP-1 cells. (A) Log_2_ fold-change and *P*_*adj*_ values of genes differentially expressed (log_2_ fold-change ≥ 2 or ≤ −2; *P*_*adj*_ < 0.01) were used in the Ingenuity Pathways Analysis^®^. Stacked bars showed pathways significantly regulated (- log(p-value) > 1.3 = *p* < 0.05; z scores > 2 or ≤ −2). Positive z scores represent pathways up-regulated and negative z scores represent pathways down-regulated by infections. (B) Overlapped pathway up-regulated (left panel) and down-regulated (right panel) canonical pathways summarized in Venn diagrams.

**Fig. 3Sb.**
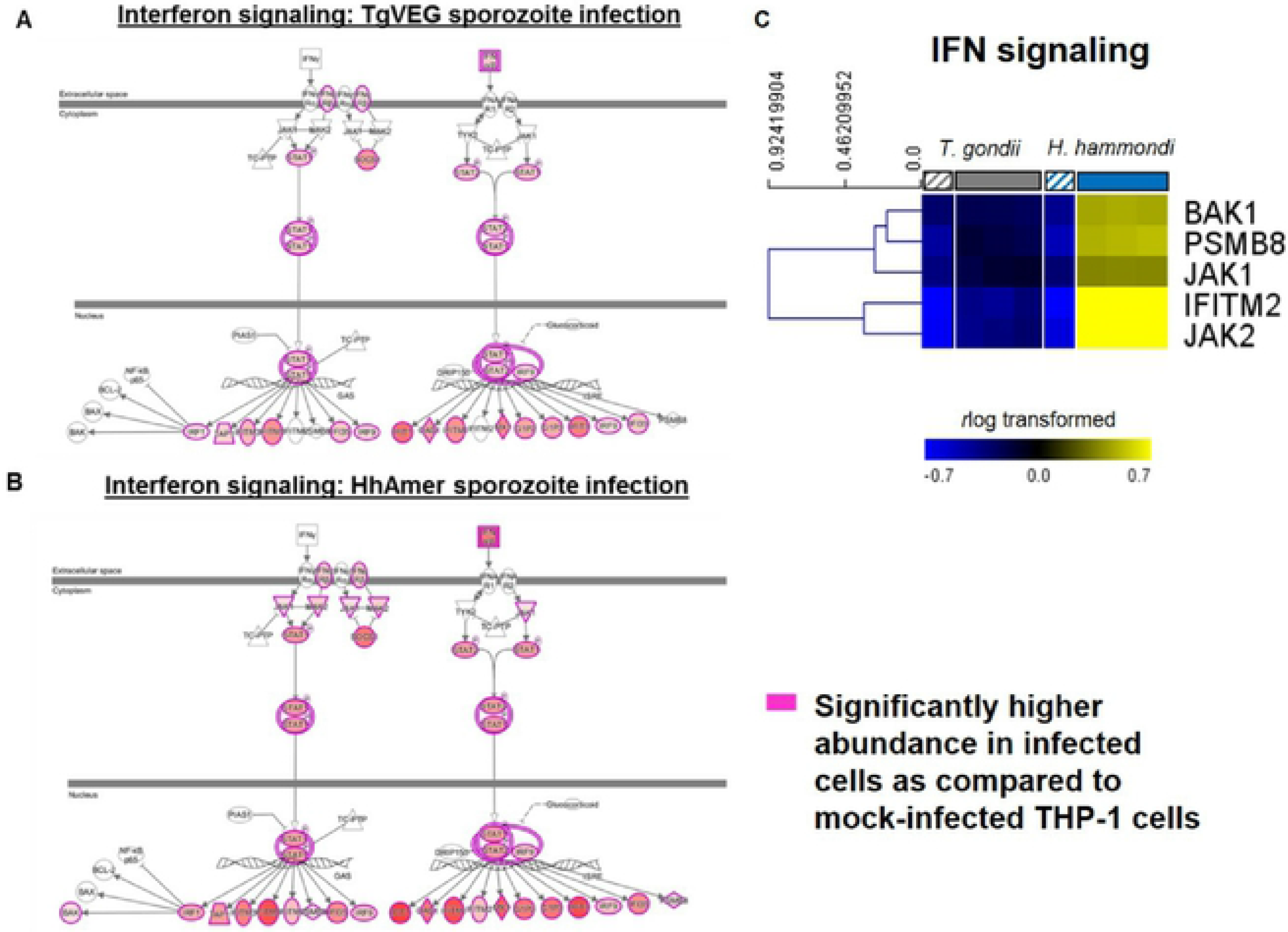
*T. gondii* and *H. hammondi-infected* cells shared similar activated pathways but to dramatically different degrees. Interferon signaling canonical pathway regulation in THP-1 cells infected with T. gondii (TgVEG; A) and H. hammondi (HhAmer; B). (C) Heatmap of gene expression (log_2_-transformed) TgVEG and HhAmer-infected THP-1 cells as in (A) and (B). Infected cells shown represented by the solid boxes whilst mock-infected cells represented by striped boxes. Data were mean-centered and hierarchically-clustered (Euclidean distance).

**Fig. 3Sc.**
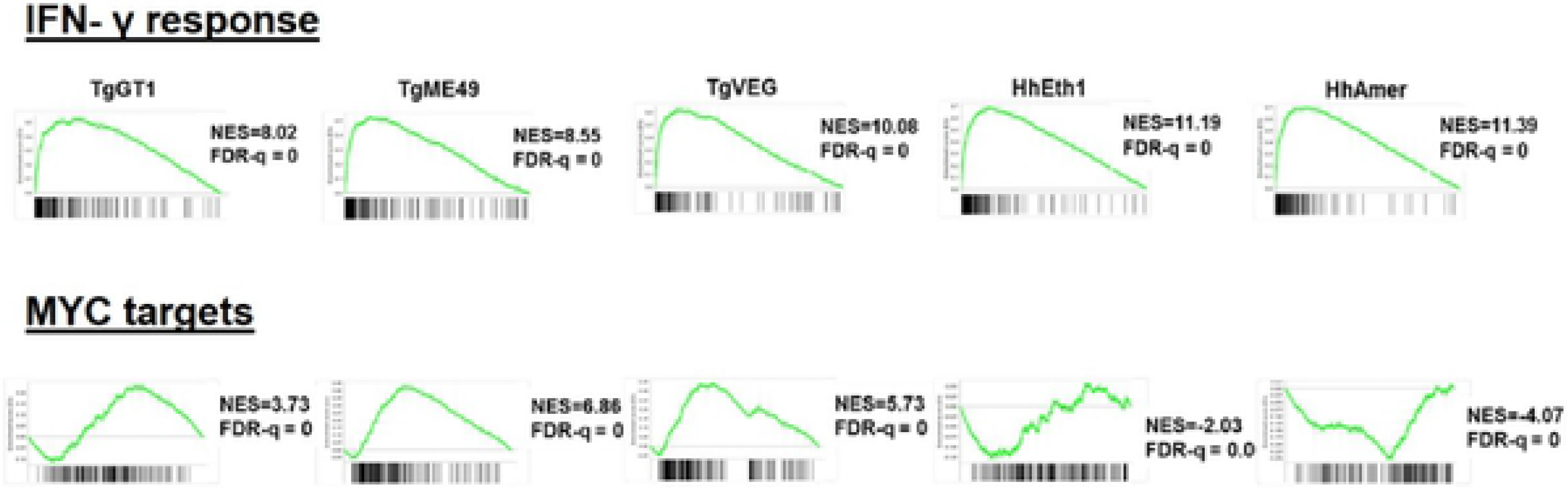
Enrichment plots of the Pre-ranked Gene Set Enrichment Analysis (GSEA) of THP-1 cells infected with *T. gondii* or *H. hammondi.* Enrichment plots from pre-ranked GSEA for *IFNy response* (top panels) and *MYC targets v1* gene sets (bottom panels) in THP-1 cells response to *T. gondii* (Tg) or *H. hammondi* (Hh) infections. Normalized enrichment scores (NES) were shown for each of the plots. *IFNy response* gene set was highly positively enriched in response to *H. hammondi* infection while MYC targets v1 gene set was negatively enriched in *H. hammondi-infected* cells as compared to *T. gondii* infections.

**Fig. 5S.**
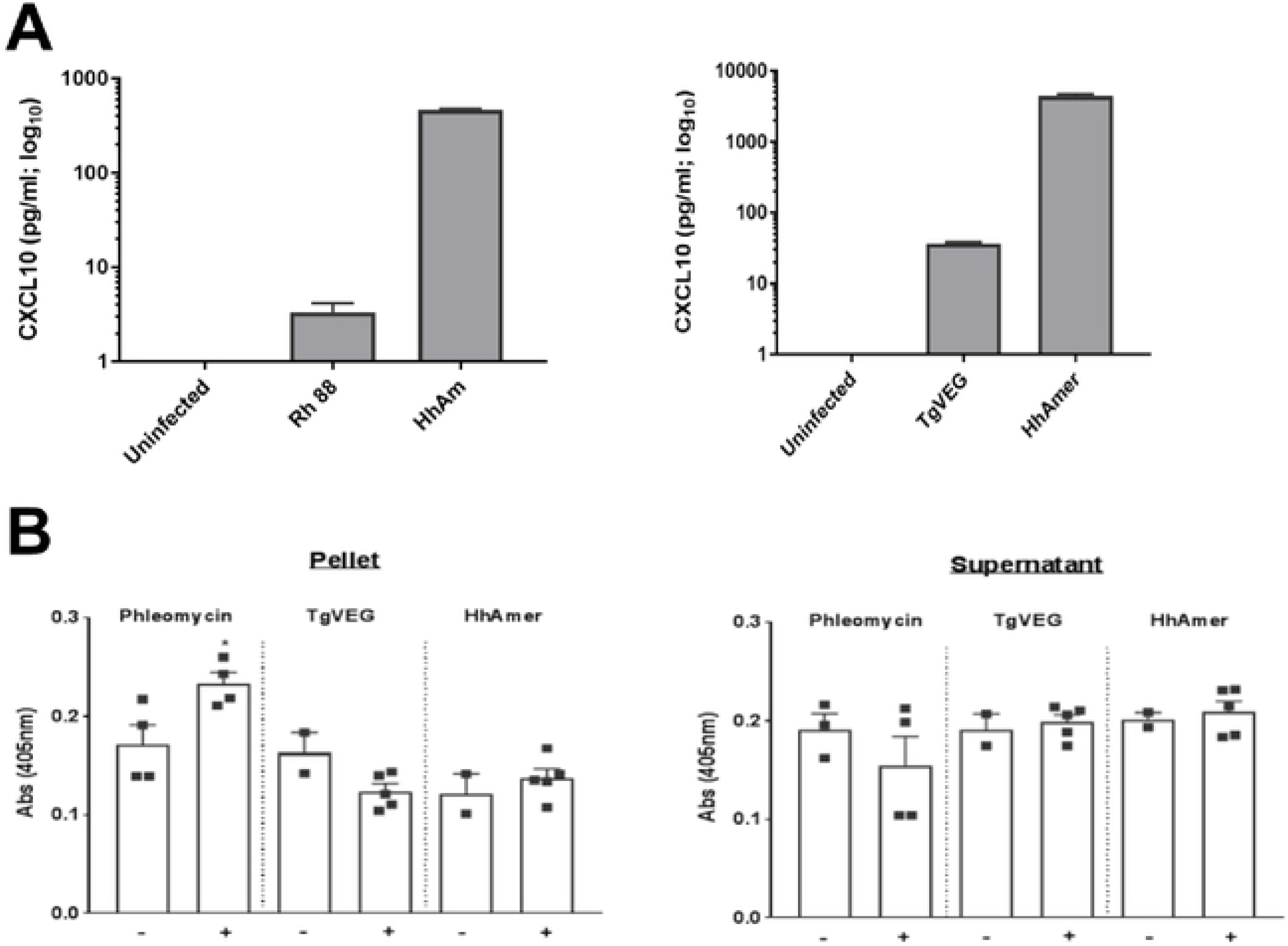
(A) CXCL10 secretion in THP-1 cells infected with *T. gondii* (Rh88 or TgVEG), *H. hammondi* (HhAmer) or uninfected THP-1 cells. THP-1 cells were infected with MOI of 2 of parasites for 20 h and supernatant was collected by pelleting the cells at 200 x g for 10 min. Secretion of CXCL10 was analyzed with ELISA. (B) β-galactosidase (gal) reagent assay to detect activity of β-gal in the THP-1 cells (Pellet) and secreted proteins (Supernatant). THP-1 cells were infected with *T. gondii* (TgVEG) or *H. hammondi* (HhAmer) with an MOI of 1.6 for 72 h and cells and supernatant were collected for β-gal activity assay. THP-1 cells were also treated with 50 µg/ul phleomycin for 72 h to induce senescence. THP-1 cells were pelleted and supernatant was collected for the assay. Cell pellet was lysed with IP lysis buffer and assayed for 13-gal activity. The 13-gal activity was significantly higher in the pellet of phleomycin-induced senescence THP-1 cells as compared to TgVEG and HhAmer infections (Tukey's multiple comparisons test, \**p*<0.05).

**Fig. 7S.**
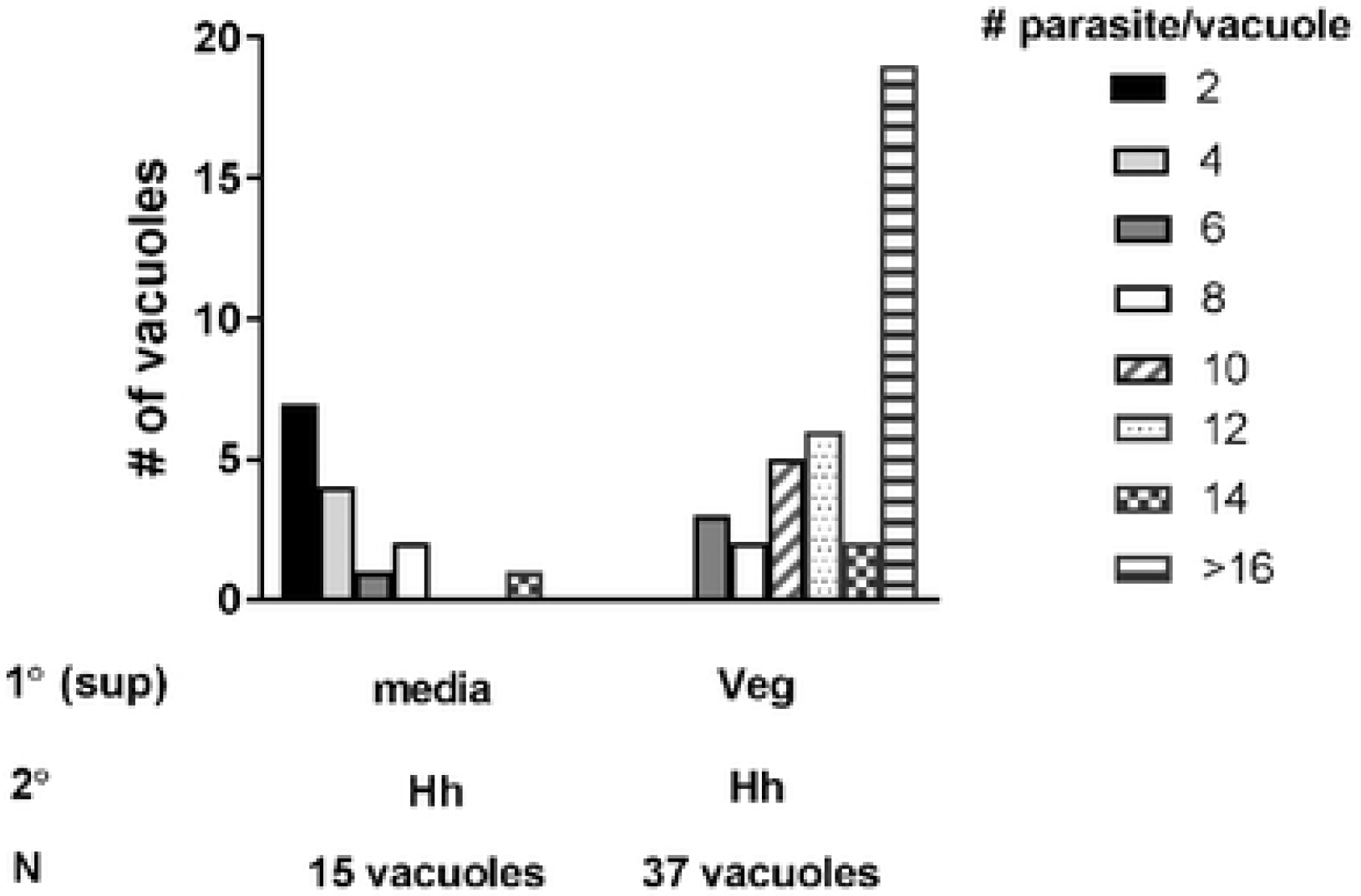
Increased *H. hammondi* vacuole size in HFF cells pre-conditioned with *T. gondii* and THP-1 cells ESP. Histogram showing vacuole size of *H. hammondi* growth in the absence and presence of *T. gondii* (Veg) and THP-1 cells ESP in human foreskin fibroblast (HFF) cells. Total numbers of vacuoles (N) were counted from one coverslip at 72 h post 2° infection. Veg and THP-1 cells ESP was prepared in THP-1 cell infected with *T gondii* for 4 h, and filtered the collected supernatant through a 0.2 µm filter. HFF cells were pre-conditioned with Veg and THP-1 cells ESP and freshly excysted Hh sporozoites were added into the same wells. Growth rates of parasite on HFF cells pre-conditioned with *T. gondii/THP-1* ESP was significantly higher than HFF cells that did not pre-condition with the ESP (****p<0.0001, unpaired *t* test comparing means of vacuole size of the coverslips).

